# EEG Cross-Frequency Phase Synchronization as an Index of Memory Matching in Visual Search

**DOI:** 10.1101/2020.09.22.306431

**Authors:** Anna Lena Biel, Tamas Minarik, Paul Sauseng

**Author notes:** Contact: Prof. Dr. Paul Sauseng, Department of Psychology, LMU Munich, Leopoldstr. 13, 80802 Munich, Germany.

## Abstract

Visual perception is influenced by our expectancies about incoming sensory information. It is assumed that mental templates of expected sensory input are created and compared to actual input, which can be matching or not. When such mental templates are held in working memory, cross-frequency phase synchronization (CFS) between theta and gamma band activity has been proposed to serve matching processes between prediction and sensation. We investigated how this is affected by the number of activated templates that could be matched by comparing conditions where participants had to keep either one or multiple templates in mind for successful visual search. We found a transient CFS between EEG theta and gamma activity in an early time window around 150ms after search display presentation, in right hemispheric parietal cortex. Our results suggest that for single template conditions, stronger transient theta-gamma CFS at posterior sites contralateral to target presentation can be observed than for multiple templates. This can be interpreted as evidence to the idea of sequential attentional templates. But mainly, it is understood in line with previous theoretical accounts strongly arguing for transient synchronization between posterior theta and gamma phase as a neural correlate of matching incoming sensory information with contents from working memory and as evidence for limitations in memory matching during multiple template search.

## Introduction

Working memory and selective attention interact in many situations of our everyday life, influencing how we perceive the world. Image yourself looking for your car keys that you must have left somewhere in the kitchen. During your search, you will scan a rich visual environment for something that matches the representation of keys that you have in mind. Such situations are commonly described as visual search. Brought to a cognitive psychology laboratory, participants in a visual search paradigm are usually asked to search for a target object among a number of distractor objects presented on a computer screen. Current theories of attention hold that when we are searching for a target, then keeping a template representation of the target in working memory – a so called attentional template – leads to a bias in the competition for neuronal resources in favour of template-matching stimuli (Bundesen, 1990; Bundesen et al., 2005; Desimone & Duncan, 1995; Duncan & Humphreys, 1989). Insight into the neural mechanisms underlying the activation of such mental templates and their comparison with sensory input comes from studies in healthy humans (Gayet et al., 2017; Soto et al., 2007; Spaak et al., 2016) patients with frontal lesions (Soto et al., 2006; Yago et al., 2004), lesion studies in primates (Everling et al., 2006; Lba & Sawaguchi, 2003; Rossi et al., 2007), as well as from formal theoretical models (Friston, 2005). Thereof, especially prefrontal brain regions are known to be involved in visual search and the top-down control of visual perception from working memory by impacting on lower visual cortex (for review, see Soto et al., 2008).

Interactions between higher and lower brain areas, as assumed to be involved in visual search, can be well investigated by analysis of oscillatory brain activity. Interaction within or between brain areas is implemented by synchronous neural activity, as reflected by rhythmical oscillations of the field potential which can be recorded using scalp electroencephalography (EEG). Oscillatory EEG activity is commonly reported to play a functional role for perceptual and cognitive processes (Buzsáki & Draguhn, 2004; Fell & Axmacher, 2011; Fries, 2005). Two brain areas are assumed to be functionally coupled when their activity is more synchronous than what would be expected from random fluctuations. It has been suggested that the complexity of the neural network(s) involved will determine the frequency range of the dynamics in a given interaction (Buschman & Miller, 2007; Fell & Axmacher, 2011; Fries, 2005), such that long-range interactions during top-down processes draw on lower frequencies in the theta band (∼6 Hz) and alpha band (∼10 Hz), whereas higher frequency, such as gamma band (>30 Hz), interactions characterize more local, small-network interactions.

Long-range interregional synchronization between human prefrontal and parietal areas has repeatedly been found for oscillatory brain activity in the theta or alpha frequency range, for example during highly demanding working memory tasks (e.g. Sarnthein et al., 1998; Sauseng et al., 2005; von Stein et al., 2000). Synchronous gamma band activity has been linked to bottom-up processes such as feature-binding and awareness (for review, see Engel & Singer, 2001), however, it also been associated with processing demands related to object representations, directed attention and active maintenance or manipulation of information (e.g. Axmacher et al., 2006; Friese et al., 2013; Jensen et al., 2007). Thereby, the common underlying mechanism is a need for comparison of sensory input with memory content as proposed by Herrmann and colleagues (2004), who suggest a central role of gamma-band responses in matching memory contents with sensory input. However, it has been argued that this model would well account for the matching with long-term memory information but less well for the matching with mental templates kept in working memory (Holz et al., 2010; Sauseng et al., 2015) so that in addition, a long-range fronto-parietal network drawing on theta band oscillations is expected to be involved.

A neural mechanism for this involvement may be phase synchronization between theta and gamma band activity, as proposed in a framework that could well account for the activation of mental templates from working memory, controlled by frontal resources and replayed into higher visual areas drawing on a theta network, and their comparison with sensory input, wherefore synchronization with gamma band phase is suggested (Sauseng et al., 2010, 2015). Theta band activity has been shown to generally have a strong influence on local cortical activity both in the human and animal brain, namely by entraining neuronal spiking and fast oscillatory activity, such as gamma band activity (Canolty et al., 2006; Fell & Axmacher, 2011; O’Keefe & Recce, 1993; Sirota et al., 2008). From studies using EEG in humans, perceptual and working memory processes have been associated with theta-gamma frequency interaction (Berger et al., 2019; Demiralp et al., 2007; Griesmayr et al., 2010; Sauseng et al., 2008, 2009; Schack et al., 2002). Such cross-frequency coupling is commonly taken as an indicator of an exchange of information between global and local neuronal networks (see Sauseng & Klimesch, 2008). Especially phase synchronization could integrate neuronal processing which is distributed into neuronal assemblies and across frequency bands by enabling consistent spike-time relationships between the oscillating neuronal populations; and cross-frequency phase-synchronized input to pyramidal layer 5 cells may facilitate neuronal bursting of these cells (Palva & Palva, 2018). So, concerning the activation of mental templates from working memory and their comparison with sensory input, cross-frequency phase synchronization between theta and gamma band oscillations can be regarded as a candidate neural mechanism underlying this process (Sauseng et al., 2010, 2015).

In the current study, we asked whether the number of activated mental templates that could be matched with sensory input does influence memory matching in visual search, as presumably reflected by a transient cross-frequency interaction between theta and gamma frequencies. Whether it is possible to look for multiple objects at the same time is a question of active debate and ongoing research. Some studies corroborate a serial bottleneck that requires alternating between items (Olivers et al., 2011; Ort et al., 2017), whereas others rather support a parallel model assuming less efficient, but parallel processing of each item (Beck et al., 2012; Hollingworth & Beck, 2016; Ort et al., 2019) or assume hybrid models (e.g. Bays & Husain, 2008). In a range of different paradigms, clear multiple target costs have been found both on the behavioral and the EEG level, indicating that multiple-target search seems to be limited in capacity, however, evaluating the exact processing stage at which serial or parallel processing limitations occur, has proven difficult or led to sometimes mixed results (for review, see Ort & Olivers, 2020). The aim of the current study was to investigate the stage of memory matching, by measuring theta-gamma phase synchronization as a proposed neural correlate.

Indeed, there is evidence in support of the involvement of a transient theta to gamma phase synchronization in posterior parietal brain areas in integrating top-down controlled mental templates with bottom-up visual processing (Holz et al., 2010; Sauseng et al., 2008). In cases where our expectancies and the actual visual input match, a higher transient phase synchronization than in case of a non-match has been found between posterior theta and gamma oscillations. For example, in a delayed-match-to-sample working memory paradigm, Holz and colleagues (2010) found stronger right-hemispheric posterior EEG theta to gamma phase synchronization for congruent in comparison to incongruent trials 150-200 ms post probe presentation. Additionally, the authors reported a resetting of theta phase shortly before this, leading them to propose that a posterior phase resetting of theta band oscillations could enable the transient cross-frequency synchronization with high frequency activity in the gamma band range found shortly after. Unexpectedly, stronger theta-high gamma phase synchronization for non-match than match was seen at left posterior recording sites. Here, Holz and co-workers speculated that this reversed effect might indicate the detection of a discrepancy between mental template and a presented item which might in turn trigger a more detailed local processing of sensory input. In the other study, however, the effect was not only right-lateralized but occurred on both the left- and right-hemispheric region of interest (Sauseng et al., 2008). Here, it was reported that in a visuospatial attention task, the increase of theta to gamma phase synchronization around 150 ms after target-onset was always larger contralateral than ipsilateral to target presentation in the validly cued hemifield. This was interpreted as a neural correlate of the matching of memory content with incoming sensory input, modulated by a top-down attentional process. This is supported by the idea that cross-frequency phase synchronization could be a candidate mechanism for integrating cognitive functions, such as the representation of sensory information and attentional or executive functions, by connecting the most central network nodes between distributed neuronal networks that support these functions (Palva & Palva, 2018).

An open question is how this proposed neural correlate of memory matching may be modulated when only one of multiple templates is met by matching sensory input. Interestingly, another form of cross-frequency interaction may also be involved during the earlier stage of the retention of multi-item working memory content. Influential computational models propose that separate memory items are represented by separate gamma waves which are nested into a theta wave (Jensen & Lisman, 1998; Lisman & Idiart, 1995) or that each item is coded by an entire gamma burst, i.e. multiple gamma waves, which are nested into a theta wave (Herman et al., 2013; Van Vugt et al., 2014). Thus, it is assumed that to hold in mind multiple templates, these need to be refreshed in a sequential manner. Although our working memory can undoubtedly represent multiple items, a prominent model proposes that the number of templates that can be active at a time is limited to only one (Olivers et al., 2011). This would predict that even though multiple templates coexist, only one of them can interact with sensory input after another. As mentioned earlier, alternatives to these serial models exist, but while parallel processing during selection and preparation may be possible, it is yet also relatively unclear whether this could generalize from paradigms with relatively simple target features (e.g. color) to paradigms utilizing more complex target stimuli (Ort & Olivers, 2020). But in any case, a single mental template should enable a fast and precise memory matching, whereas a larger number of mental templates that could potentially be matched to visual input should come at costs that disable such an early and precise matching process. Thus, visual search for multiple templates can be expected to come along with limitations in the memory matching stage, whether serial or parallel in nature. These limitations should be reflected in a transient theta to gamma phase synchronization in posterior parietal brain areas, if this mechanism is indeed involved in integrating top-down controlled mental templates with bottom-up visual processing, as proposed (Sauseng et al., 2010, 2015).

More specifically, on the basis of these abovementioned models, the proposed neural correlate of memory matching should be modulated when only one of multiple templates could be matched to sensory input in the following way: Assuming that we keep multiple templates in mind sequentially, one would assume that upon search display presentation in a given trial, the first, second or n^th^ item in the sequence incidentally matches sensory input. Further assuming that only one of them can interact with sensory input, memory matching should occur relatively early, a bit later or even much later in a given trial, depending on whether the sequence’s first, second, or a later mental template could be matched to the current visual input. This means that in conditions where multiple mental templates could be matched to one out of several possible targets appearing on screen, the memory matching mechanism and likewise its neural correlate, is supposed to display more temporal variability across trials. Therefore, lower overall theta to gamma phase synchronization values, which are measured through an index aggregated over trials, are expected than when a single mental template enables a precise matching and thus a temporally aligned theta to gamma phase synchronization is expected.

A sequential matching process would be a rather plausible interpretation of low estimates of cross-frequency phase synchrony in multiple template search. However, if memory matching in a multiple template search happened with great temporal variability and also consistently later than in a single template search, or if there was more temporal variability in theta-gamma phase relations due to other unspecific differences imposed by multiple template search, then low phase synchronisation estimates would be expected as well. Low phase synchronisation estimates would also be expected if memory matching did not take place at all during multiple template search; however, this would really only be plausible when none out of multiple templates can be matched, such as previously found in non-match trials from other task paradigms (Holz et al., 2010; Sauseng et al., 2008, 2009). Conversely, when assuming that multiple templates can be matched in parallel, but with costs due to mutual competition, then slightly delayed, but high estimates of phase synchrony similar to a single template search would be expected. In any case, limitations in memory matching due to multiple template search should be reflected in a transient theta to gamma phase synchronization in posterior parietal brain areas, if assuming that this mechanism is indeed a neural correlate of memory matching. Not all of these options can be disentangled due to the nature of the theta-gamma phase coupling index, but in any case, if a modulation of the transient cross-frequency interactions between theta and gamma frequencies was observed during multiple template search compared to single template search, this would indicate that the number of activated mental templates that could be matched with sensory input does influence memory matching in visual search.

We designed a visual search paradigm where displays with four abstract symbols were shown to participants, each display containing one target among distractors. Participants had to indicate in which quadrant of the display their target symbol had been presented. We varied the number of mental templates that had to be kept in mind for successfully performing the visual search. In separate experimental blocks, the target could be either one single symbol (i.e. one item had to be held in memory) or one out of a set of three target symbols (i.e. three items would have to be retained). In the single template condition, we expected that around 150-200 ms after search array onset, a transient increase in theta-gamma phase synchronisation should arise over right-hemispheric posterior brain areas for targets located in the contralateral hemifield, relative to ipsilateral targets, because this would corroborate evidence from other task paradigms (Holz et al., 2010; Sauseng et al., 2008, 2009). Conversely, such transient increase in phase synchronisation should not arise in a condition where three mental templates were required for successful search performance, because a larger number of mental templates that could potentially be matched to visual input would modulate cross-frequency phase interactions.

## Methods

### Participants

Thirty-five typically developed volunteers participated in the experiment. All gave written informed consent prior to their participation and received financial compensation or course credits upon completion. Four participants had to be excluded from analysis because their percentages of correct responses were in the range of chance level, suggesting they were merely guessing, in at least one condition of interest. Two more participants were excluded based on too noisy EEG recordings. In the remaining sample that was included in the analyses (n=29), mean age was 24.7 years (*SD* = 2.8) and 7 participants were male. All but one were right-handed, as assessed by the Edinburgh Handedness Inventory (Oldfield, 1971) and all reported normal or corrected to normal vision. The study was approved by the local Ethics Review Board and conducted according to the Declaration of Helsinki.

### Apparatus

Participants were seated in a comfortable chair in a dimly lit room and were wearing an EEG cap (Easycap®) for registering EEG signals. They had a standard computer keyboard placed on their lap. Their left and right index and middle fingers were placed on four buttons of the numbers block, namely buttons 1, 2, 4, and 5, which were marked by coloured stickers. Each button represented one of four quadrants of a visual search display. Stimuli were presented on a 22-inch Samsung S22C450 monitor with a resolution of 1280 x 1024 and a 75 Hz refresh rate, which was placed centrally and at a distance of 80 cm from an observer. Stimulus presentation was controlled using Presentation 0.71 (Neurobehavioural Systems®), which was synchronized with recording of the EEG signals in BrainVision Recorder 2.0.4 (BrainProducts®).

### Task

We recorded EEG from participants while they completed a visual search task, where they searched for a target stimulus among distractors. At the beginning of each trial (see figure 1A), a central fixation cross was presented for a random duration between 600 and 1000ms, which participants were instructed to fixate during the whole trial. Next, the search display was displayed for a duration of 200ms, and immediately masked for 1000ms. The fixation cross remained on the screen for another 1500ms. The target stimulus was presented equally often in each quadrant of the search display (25% of the trials). Participants indicated in which quadrant of the search display the target stimulus had been presented by pressing the respective button on the numbers block, for upper left (button 4) upper right (button 5), lower left (button 1) or lower right (button 2). They were instructed to respond as accurately as possible, and as soon as possible after presentation of the search display. So accuracy was emphasized over speed. As an inter-trial interval, a blank screen was shown for a random duration of 800 to 1200ms, adding up to a total trial duration of 4500ms, before the next trial started. Participants were instructed to keep their eyes fixated at the central fixation cross during the whole task.

**Figure 1.**
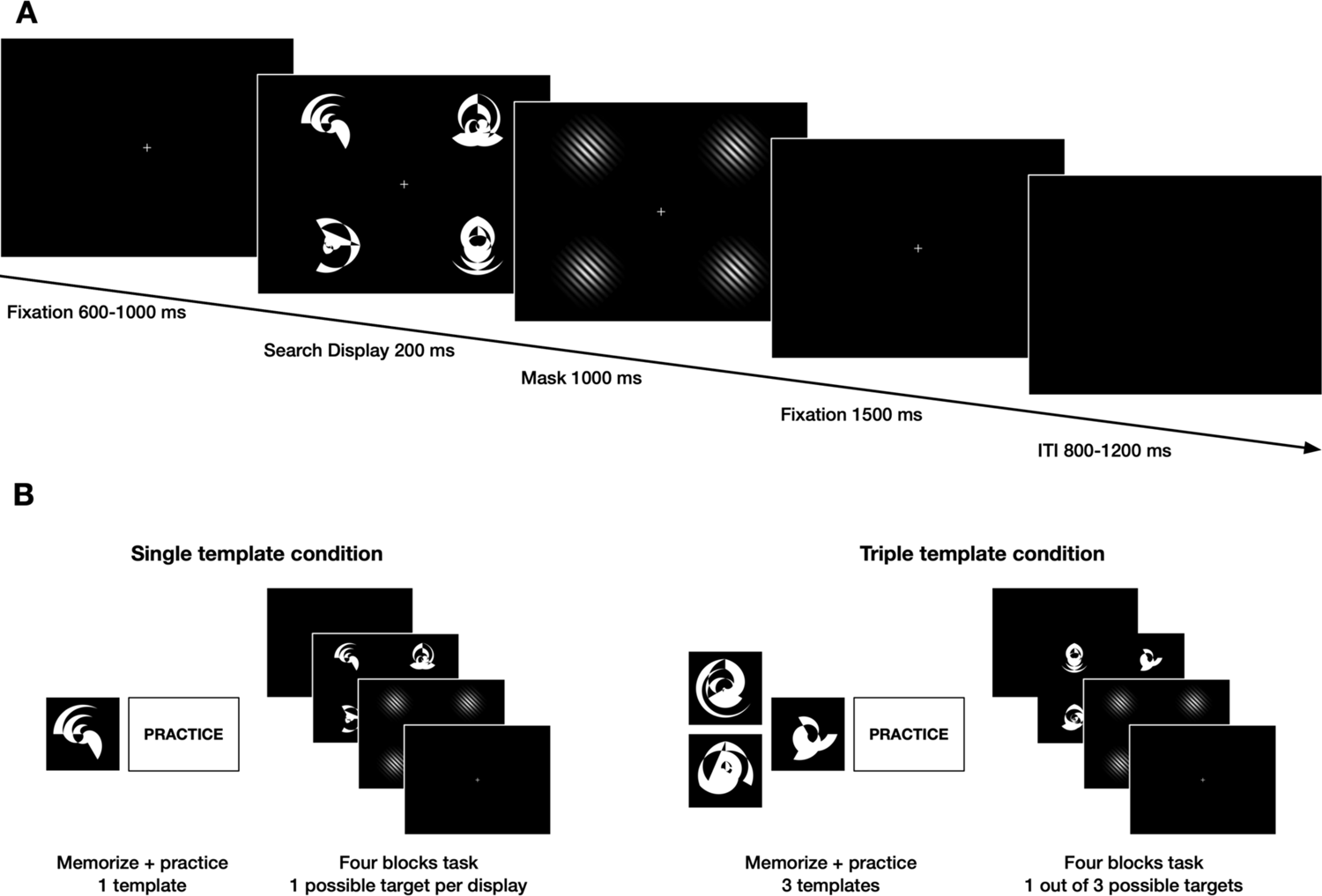
Illustration of the visual search task. A: Exemplary trial sequence of the visual search task. Trials consisted of a fixation (600-1000ms), a search display (200ms), a mask screen (1000ms), and a fixation (1500ms); Inter-trial intervals (ITI) showed a blank screen (800-1200ms). Search displays contained one target among three distractors and participants indicated in which quadrant of the search display the target had been presented by button press. B: Experimental procedure. The experiment comprised two parts with counterbalanced order across the four experimental versions (shown here: version IV). Prior to each part, consisting of four task blocks each, participants memorized and practiced the target(s).

### Stimuli

All stimuli were presented against black background, with a white fixation cross in the centre of the screen (see figure 1A). As stimuli, 16 different abstract symbols were created (for code and stimuli, see https://osf.io/wyezs/). None of these abstract symbols were known to the observers. Thus, participants could not rely on existing semantic memory contents such as the name of the target feature and we assumed that the search for this kind of complex targets would have relied more on an active attentional template of the visual target(s) in working memory. Four stimuli were used as targets (1-4) and 12 other stimuli as distractors (A-L).

Our paradigm contained two conditions (*Template*; single vs. triple), in which either one or one out of three possible targets was presented among distractors. To counterbalance which target and distractor stimuli appeared in the single vs. triple template conditions, four experimental versions were used (*Version I:*1 and A-F vs. 2, 3, 4 and G-L; *Version II*: 2 and G-L vs. 1, 3, 4 and A-F; *Version III*: 3 and A-F vs. 1, 2, 4 and G-L; *Version IV*: 4 and G-L vs. 1, 2, 3 and A-F). For each template condition, target and distractor stimuli were composed into 48 different search displays. Thus, in the *single template condition*, 12 search displays had the target symbol in the same quadrant, while three distractor symbols, randomly drawn from a subset of 6, were placed into the remaining three quadrants. For the *triple template condition*, a display could contain one out of three possible targets, thus each target was placed in each quadrant four times, accompanied by three randomly drawn distractors. This resulted in a total of 256 search displays being used. Mask screens displayed circular Gabor gratings at the same four locations of the search items, consisting of 9 white and 10 black lines each and oriented vertically (135°).

### Procedure

Our conditions in which the number of possible targets, and thus the number of mental templates to be held in mind, could be either one or three (*Template*; single vs. triple) each consisted of 192 trials, divided into four blocks with 48 trials, such that the experiment comprised eight blocks in total. Whether participants started with the four blocks of the single or of the triple template conditions was counterbalanced across different versions of the experiment (Version I and III: triple, then single; Version II and IV: single, then triple). Version was randomly assigned to a participant and also determined which targets and distractor sets were assigned to which condition (see apparatus and stimuli). The search displays in the single template condition always contained the same target among distractors, whereas and in the triple template condition, one out of three possible targets was presented among distractors for search. In the triple template condition, there are trials where a different or the same target as on the previous trial is presented, however, there is a much lower number of stay trials than switch trials because the paradigm was not designed to contrast these. While this does not leave us with a sufficient number of trials to analyse potential target switch costs after, we provide an overview about potential hypotheses for future studies investigating these in the supplemental materials S9. Within a block, search displays were presented in randomized order. In the triple template condition, targets appeared equally often. Participants took breaks between blocks, resulting in a total of about 40 minutes to complete the experiment.

In order to familiarize the participants with the targets, a training was completed prior to each condition (see figure 1B). During a memorization phase before each practice block, the respective single target or three targets were shown to participants in a printed version, and participants were asked to memorize them well. Practice blocks had fewer trials than the actual experimental blocks and served to make participants familiar with their targets. The same target(s) and distractors as in the respective template condition were displayed here, however, only with one instead of four stimuli on screen. Upon target presentation, participants decided whether the displayed stimulus was (one of) their target(s) or not. They did so with a button press, where they were asked to press left (no target) and right (target) arrow buttons on the keyboard. At least two memorization phases and practice blocks were completed per condition. If necessary, more were administered, until participants were confident in discriminating between target(s) and distractors and performed well above chance in doing so. Note that the training beforehand was necessary because the target stimuli in our task were displayed briefly and were rather complex (for details, please see below) and unknown to the observers. We intended to build a task that was effortful and where participants could not rely on existing semantic memory contents such as the name of the target feature. We assumed that the search for this kind of complex targets would have relied more on an active attentional template in working memory (Gunseli et al., 2014). For this effortful search, though, it was not possible to ask participants to memorize a trial-by-trial changing target, because they could not rely on one distinct feature, but instead the abstract figure as a whole. Therefore, we kept the target(s) constant in each condition and trained participants beforehand. This makes it possible that participants may have stored those memorized target(s) in long-term memory before the start of our task. We elaborate on this in the discussion.

The whole experiment, including the preparation of the EEG cap, instructions, breaks, training blocks and experimental blocks, took about 2 hours.

### EEG Data Acquisition and Preprocessing

EEG was registered from 60 scalp locations with Ag-AgCl electrodes arranged according to the extended 10-10-system in a TMS compatible electrode cap (Easycap®), using a BrainAmp MRplus amplifier (BrainProducts®). Two electrodes were placed above and next to the left eye for recording horizontal and vertical eye movements and blinks. An additional ring-electrode on the tip of the nose was used as a recording reference and the ground electrode was placed at electrode position FPz. Electrode impedances were kept below 15 kΩ. EEG data were digitized at 1000 Hz in a frequency range above 0.016 Hz. A notch filter was set at 50 Hz. Butterworth zero phase filters were used.

EEG data were pre-processed using BrainVision Analyzer 2.0.4 (Brain Products®). Raw data was re-referenced using an average reference of all EEG channels. After filtering with a low-cutoff of 0.5 Hz (48 dB/oct) and a high-cutoff of 100 Hz (48 dB/oct), visual inspection was used for excluding data sections with large artifacts during task breaks. Next, semiautomatic Ocular Correction with Independent Component Analysis (Ocular Correction ICA) was applied to correct for artifacts caused by eye blinks and eye movements. Only trials including a correct response that was given within 3000 ms after search display onset were retained. Data were then segmented into 2000ms epochs to avoid edge artifacts in later analysis steps, ranging from 1000ms before to 1000ms after onset of a search display. Finally, epochs that contained remaining artifacts due to eye movements or muscle activity were rejected manually. On average, the number of trials that remained after these procedures were 75.2 trials (78.3%) for targets on the left side of the screen and 79.3 trials (82.6%) for targets on the right in the Single template condition. In the Triple template condition, on average 54.0 trials (56.2%) remained for left hemifield target positions and 55.2 trials (57.5%) for targets on the right.

### Cross-frequency phase synchronization index

Source-space EEG signals obtained from the brain regions of interest (ROIs; see next section for details) were decomposed using continuous wavelet transformation using Morlet wavelets. In order to extract one lower frequency band that is centred over the typical theta range and comparable to the study by Holz and colleagues (2010), for several lower frequency bands, wavelet coefficients were extracted with 5 frequency steps ranging from 1 Hz to 12 Hz, using a 5-cycle complex Morlet parameter. Thus, the frequency of interest for theta band activity had a central frequency of 6.50 Hz (bandwith 5.2–7.80 Hz). Both the theta and gamma band activity was derived using the same Morlet parameter, to be able to detect a transient modulation of phase in the gamma frequency range. Activity in several higher frequency bands was extracted with 6 frequency steps ranging from 30 Hz to 80 Hz and with a 5-cycle complex Morlet parameter. The purpose of this was to extract gamma bands that are comparable to the study by Holz and colleagues (2010) and to cover the whole range from 30 Hz to 80 Hz, but with little overlap to avoid redundancy and to reduce data for statistical analysis. Thus, three of these higher frequency bands were extracted as frequencies of interest for gamma band activity which were centered around 40 Hz (bandwith 32–48 Hz), 60 Hz (bandwith 48–72 Hz), and 70 Hz (bandwith 56–84 Hz).

Next, continuous phase values were extracted from the wavelet coefficients’ complex values, for lower and higher frequency bands. To quantify their phase consistency across trials, we calculated the cross-frequency phase synchronization index (PSI), similar to Schack and Weiss (2005) or Palva and colleagues (2005), through custom-made scripts in MATLAB R2015b. So first, for each trial and sampling point, slow frequency band and high frequency band phase values were multiplied with the central frequency of the other band. Next, the phase differences across these adjusted signals was calculated for each trial and sampling point by subtracting sampling point-wise high frequency from low frequency adjusted signals. This generalized phase difference is described with the equation (where m and n are the central frequencies of the low and high frequency bands, which are multiplied with the instantaneous phase values in the kth trial and at sampling point t for the low and high frequency f_m_ and f_n_, respectively): 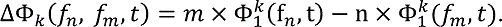 PSIs across trials were calculated as the average vector length of these generalized phase differences, by taking the square root of the sum of the squared sine and cosine values of the phase differences, averaged across trials, yielding an index ranging from 0 to 1, where 1 indicates largest synchrony in phase. From these sampling point-wise PSIs, we then created averaged PSIs for time windows of 50ms length, starting at stimulus onset up to 450 ms and an averaged PSI for a pre-stimulus time window of 200 ms, starting 200ms pre-stimulus up to stimulus onset. These were transformed using Rayleigh’s *Z (rzPSI = n × PSI^2^)*. This was done to account for the number of trials (n) that went into calculation of the index which were overall lower in the triple template condition (only correct, artifact-free trials were entered into the index, see above), since usually, measures of phase-synchronization are sensitive to the difference of trial numbers across conditions used (Cohen, 2014). Note that this yields an index not ranging from 0 to 1, but ranging from 0 to n, where larger values indicate larger synchrony in phase.

### Regions of interest and time of interest

Brain regions of interest (ROIs) for analysis of posterior theta and gamma band activity were identified in source space in order to attenuate effects of volume conduction and to reduce multi-channel EEG data. We therefore transformed EEG data from scalp-level data into voxel-based Low Resolution Electromagnetic Tomography (LORETA) data (Pascual-Marqui et al., 2002) using BrainVision Analyzer 2.0.4 (Brain Products®). Here, a standard brain based on the MNI-305 brain template and a 3-shell spherical head model is used, and the source space comprises the cortical gray matter and hippocampus in the Talairach atlas with 2394 voxels at 7 mm spatial resolution. Based on the literature (Holz et al., 2010; as well as Sauseng et al., 2008), we were interested to compare posterior theta and gamma activity in bilateral posterior ROIs. While we do not assume that these are the only brain areas involved in this task, a source-specific analysis in the study by Sauseng and colleagues (2008) showed strong effects of cross-frequency phase synchronization in a similar task, with bilateral posterior sources located within extrastriate areas, covering the left and right superior occipital gyrus for the majority of subjects. Thus, for all further analyses in source space, we manually selected bilateral posterior ROIs in the left and right superior occipital gyrus (see figure 2 for a visualization of the left and right superior occipital gyrus, as implemented in the AAL atlas (Tzourio-Mazoyer et al., 2002)): Based on MNI coordinates, we selected two LORETA voxels which are located centrally within superior occipital gyrus for a left superior occipital ROI (centroid at MNI: −15, −95, 15) and the homologue right superior occipital ROI (centroid at MNI: 15, −95, 15). For the source-space EEG signals obtained from the ROIs, we computed cross-frequency phase-synchronization indices (for details, see previous section).

**Figure 2.**
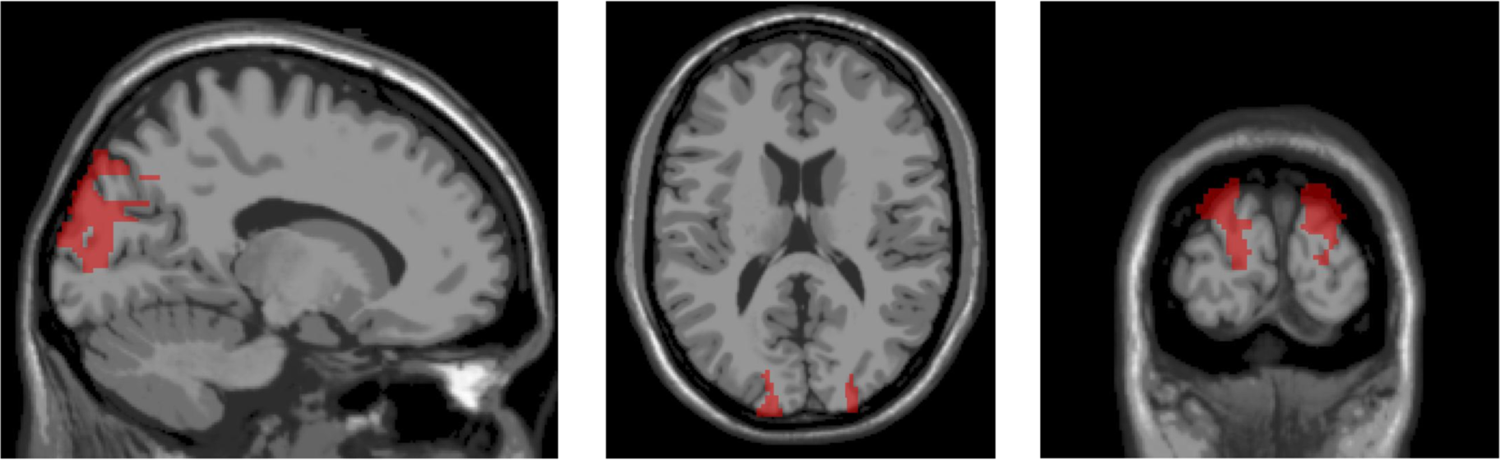
Illustration of the selected regions of interest (ROIs) in source space. A left middle occipital ROI (centroid at MNI −15, −95, 15) and the homologue right middle occipital ROI (centroid at MNI 15, −95, 15) were selected for all further analyses in source space. Highlighted in red are the left middle occipital gyrus and right middle occipital gyrus.

Within these ROIs, we analysed differences between our experimental conditions in the time window of interest. The time window of interest (TOI) was 150-200 ms after visual search display onset, a typical time window found in the previous studies (Holz et al., 2010; as well as Sauseng et al., 2008).

### Event-related Potentials

To relate our data to the existing EEG research on visual search, in which the N2pc ERP component has been described as an important neural signature (Eimer, 2014; Luck, 2012), we computed scalp-level grand average ERP waveforms for left and right target locations in both template conditions. These were filtered between 0.5 Hz (48 dB/oct) and 35 Hz (48 dB/oct) and baseline-corrected using a pre-stimulus time window of 200 ms, starting 200ms pre-stimulus up to stimulus onset. ERPs were averaged for posterior parietal electrodes PO7 and PO8 contra- or ipsilateral relative to target location. Based on visual inspection (see figure 3), ERP waves began to differ between contralateral and ipsilateral target presentations from 220 ms onwards, which is in the N2pc latency range. The N2pc, which is consistently found in visual search tasks, is an ERP component exhibiting an enhanced negativity at posterior electrodes contralateral to target presentation and is typically interpreted as an electrophysiological marker of attentional capture. We computed the average N2pc amplitude in the time window 200-350 ms for the difference between contra minus ipsilateral sites relative to target location. Average N2pc amplitudes significantly differed between the template conditions (t(28)=-3.6, p=0.001, paired-samples t-test).

**Figure 3.**
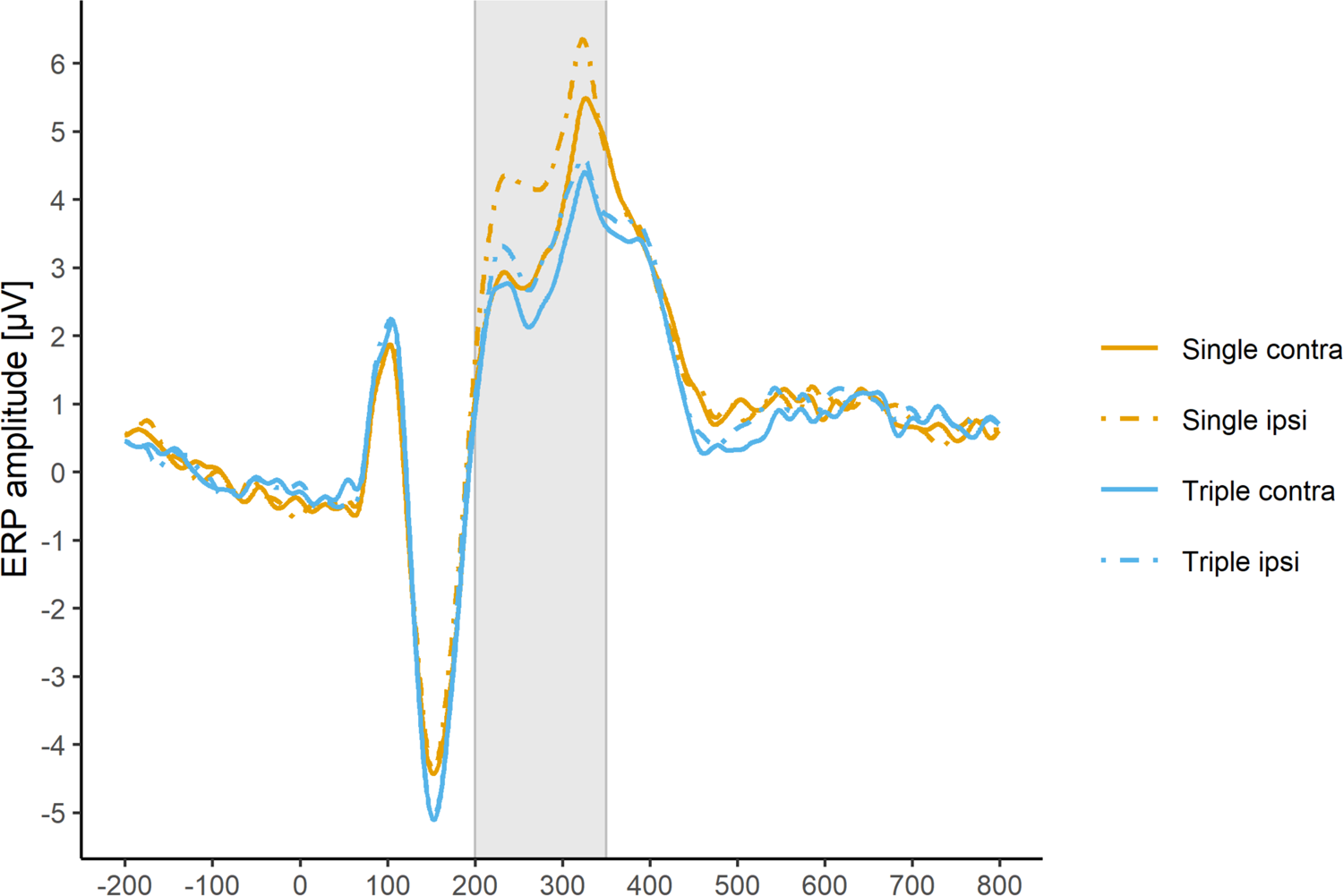
Event-related potentials (ERPs) averaged for posterior parietal electrodes PO7 and PO8, contra- or ipsilateral relative to target location in both template conditions. The time window for which N2pc amplitudes were computed is illustrated in grey.

### Behavioural data

As a measure of task performance, the percentage of correct responses was computed for each subject and in both template conditions. Additionally, the median across reaction times from trials with a correct response was calculated (note, however, that the task instructions had emphasized accuracy over speed, so reaction times should be interpreted with this in mind).

### Statistical methods

For both the behavioural and the EEG data, statistical analyses were carried out using statistical software R 3.6.1 (R Core Team, 2019) and for data visualization, plots were created using the package ggplot2 3.2.1 (Wickham, 2016). To compare behavioural data from the single template condition vs. triple template condition, two-tailed paired-samples t-tests were used on task accuracy and response times. For the analysis of EEG data, linear mixed effects models (LMMs) were implemented with the lme4 package 1.1.21 (Bates et al., 2007), contrasts matrices were derived using the hypr package 0.1.7 (Rabe et al., 2020; Schad et al., 2019) and model summary tables were produced using the lmerOut package 0.5 (Alday, 2018). As an advantage over traditional repeated-measures ANOVA, LMMs estimate the difference between conditions directly and without the need for post hoc tests instead of only the significance of a difference between conditions. Using LMMs allowed us to model random effects by subject (but as we analyze an aggregated index across trials, we could not include random effects by item as well). They also accommodate shrinkage, such that extreme and therefore less reliable estimates from individual subjects are shrunk towards the grand mean, producing more reliable estimators (see Gelman & Hill (2007) or Pinheiro & Bates (2000) for a general introduction into mixed regression models and Payne et al. (2015) and Alday et al. (2017) for an overview on LMMs, parameter estimation and model fitting, examples of their use for EEG data analyses and further literature recommendations on LMMs). The consistency of theta-gamma phase difference, as measured by the Rayleigh’s z-transformed cross-frequency phase synchronization indices (rzPSIs), was analysed for condition differences separately for the left and right posterior ROI (see above for details on the ROIs). This was done because data stemmed from sources located in separate hemispheres and because other studies from our group have previously found lateralized effects of cross-frequency synchronization (see Sauseng et al. (2009) or Holz et al. (2010)).

While the whole time series of rzPSIs was inspected descriptively, data were analysed for condition differences exclusively in the time window of interest found in previous studies, 150-200 ms after target onset (see above for details on the TOI) to reduce data for statistical analysis. Here, based on our hypothesis, we were mainly interested in an interaction between template condition and target location or any higher-order interaction involving these two factors. Adding rzPSIs for theta-to-40 Hz, theta-to-60 Hz, theta-to-70 Hz into the model enabled us to assess whether a broadband or rather frequency specific theta-to-gamma band effect were involved. In separate analyses for each ROI, we used a LMM where rzPSIs from that ROI were predicted by the fixed effects COND (Template condition: single, triple), TARG (Target location: contralateral, ipsilateral), CFS (Cross-frequency synchronization: theta-to-40 Hz, theta-to-60 Hz, theta-to-70 Hz), and their interactions. The model included a single random-effects term for the intercept of the individual subjects SUBJ. For LMM modeling, the categorical variables were encoded with sequential difference contrasts (for 2-level predictors COND and TARG: (1/2, - 1/2); for 3-level predictor CFS: (−2/3, 1/3, 1/3) and (−1/3, −1/3, 2/3)). Thus, the intercept is estimated as the grand average across all conditions and resulting fixed effect estimates can be interpreted as main effects. For the model summaries we regarded contrast coefficients with absolute *t* values larger than 1.96 as indicative of a precise estimate. T-values above 1.96 can be treated as approximating the two-tailed 5% significance level since a *t*-distribution with a high degree of freedom approaches the z distribution (Baayen et al., 2008). The reported models were fit based on restricted maximum likelihood estimation.

### Data and code availability

Data and code needed to reproduce all reported findings are available in our data repository (https://osf.io/h2j6d/).

## Results

### Behavioural analyses

Task accuracy (measured as the percentage of correct responses) was higher in the SINGLE template condition (M = 86.08%, SD = 14.51%) than in the TRIPLE template condition (M = 63.031%, SD = 11.56%) as indicated by a significant paired samples t-test (t(28) = 5.94, p<0.001, d = 1.76). Similarly, reaction times (computed as the median across correct trial’s reaction times) in the SINGLE template condition (M = 698.03ms, SD = 276.14ms) were significantly faster than in the TRIPLE template condition (M = 888.07ms, SD = 322.22ms; t(28) = −3.87, p < 0.001, d = −0.63). Both these results indicate that behavioural task performance was better when participants had to search for one target among distractors than for one out of three possible targets.

### EEG analyses

#### Theta-gamma phase synchronization in the right hemispheric ROI

Figure 5 shows single-subject rzPSIs and their group average from the right hemispheric ROI in the time window 150-200 ms after visual search display onset. A summary of model fit for rzPSIs from the right hemispheric ROI in the time window 150-200 ms after visual search display onset and the fixed effects COND (single, triple), TARG (contralateral, ipsilateral), CFS (theta-to-40 Hz, theta-to-60 Hz, theta-to-70 Hz), their interactions as well as the random effect for SUBJ can be seen in table S1.1 in the supplemental materials S1. A visualization of the fixed effects is provided in figure 4A. The grand mean rzPSIs have an estimate of 0.8 as represented by the intercept. TARG has an effect (0.062, t = 2.5) indicating that targets located in the contralateral hemifield elicited larger rzPSIs than targets at ipsilateral locations, but there also is an interaction between COND and TARG (0.18, t = 3.6), indicating that this target-related difference is larger in the single than in the triple template condition (see figure 4B). Importantly, no other contrast involving the interaction effect between COND and TARG exceeded the threshold of absolute t values larger than 1.96. So this critical effect does not interact with gamma frequency dependent differences, although three contrasts involving the factor CFS yield precise estimates with absolute *t* values larger than 1.96: One contrast shows that rzPSIS for Theta-to-70Hz are smaller than for Theta-to-60Hz (−0.068, t = −2.2), however, there also is an interaction with COND, reflecting that this gamma frequency dependent difference in rzPSIs is smaller for single than triple template conditions (0.15, t = 2.5). The interaction between the other CFS contrast and COND indicates that the difference between rzPSIs for Theta-to-60Hz and for Theta-to-40Hz is larger for single than triple template conditions (−0.14, t = −2.3). Essentially, both these interaction effects involving the factor CFS are driven from overall smaller rzPSI estimates in the single compared to the triple condition for Theta-to-60Hz, whereas the two template conditions have similar RzPSI estimates for Theta-to-40 Hz and Theta-to-70 Hz (see figure 4C).

**Figure 4.**
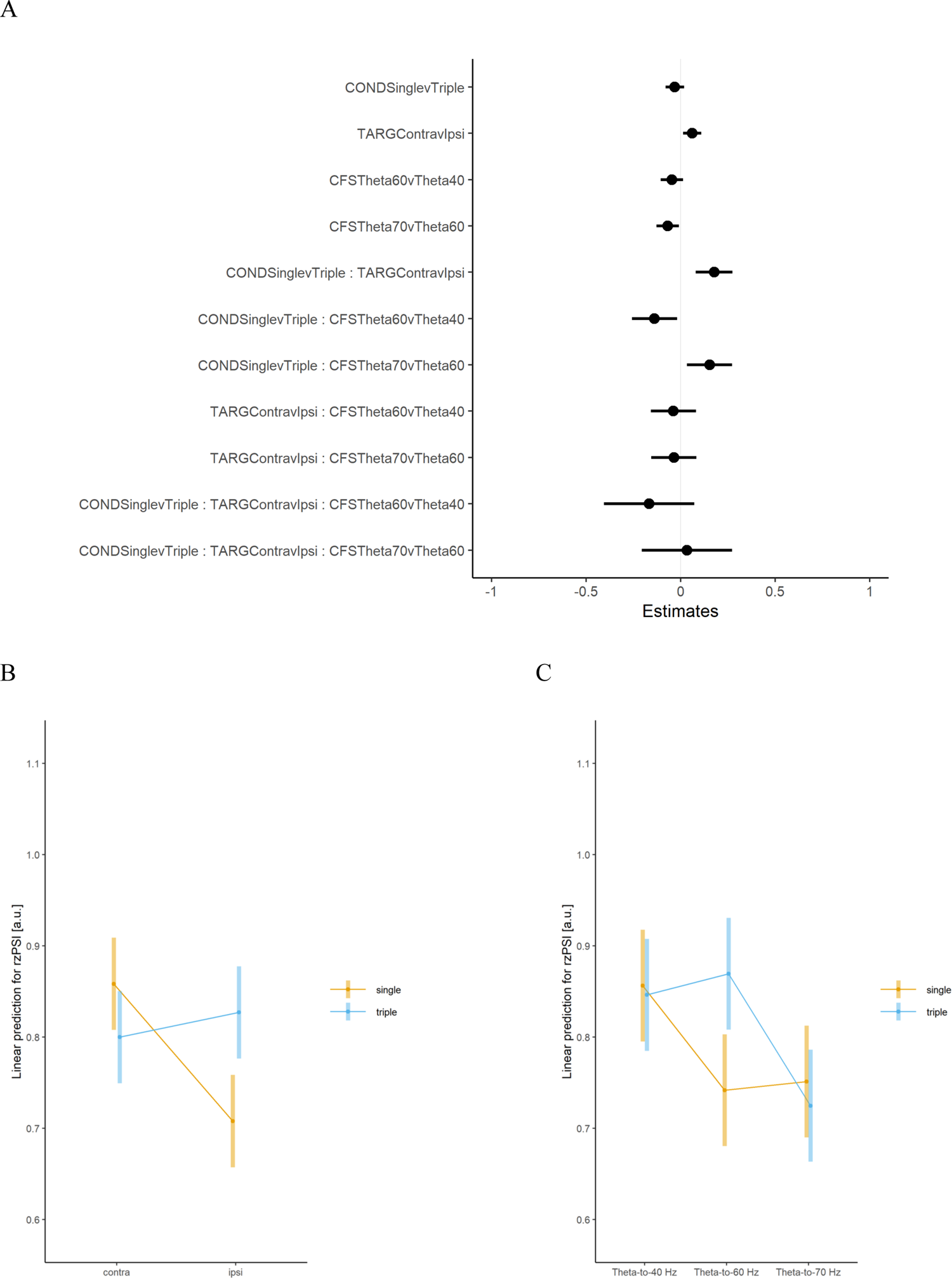
Visualization of fixed-effect estimates for model fit (A) of the cross-frequency phase synchronization indices (Rayleigh’s z-transformed; rzPSIs), measuring the consistency of theta-gamma phase difference, from the right hemispheric ROI in the time window 150-200 ms after visual search display onset. The model includes a random-effects term for the intercept of individual subjects and the fixed effects COND (single, triple), TARG (contralateral, ipsilateral), CFS (Theta-to-40 Hz, Theta-to-60 Hz, Theta-to-70 Hz), and interactions between them.Linear prediction for rzPSIS from the model showing the substantial effects for the interaction contrast between COND and TARG (B) and the interaction contrasts between COND and CFS (C). The substantial main effect TARG is not shown separately due to its involvement in the interaction with COND. Note: Dots represent values of the estimated coefficients and lines show their standard deviations in panel A. In panels B and C, dots represent estimated marginal means and lines their confidence intervals.

**Figure 5.**
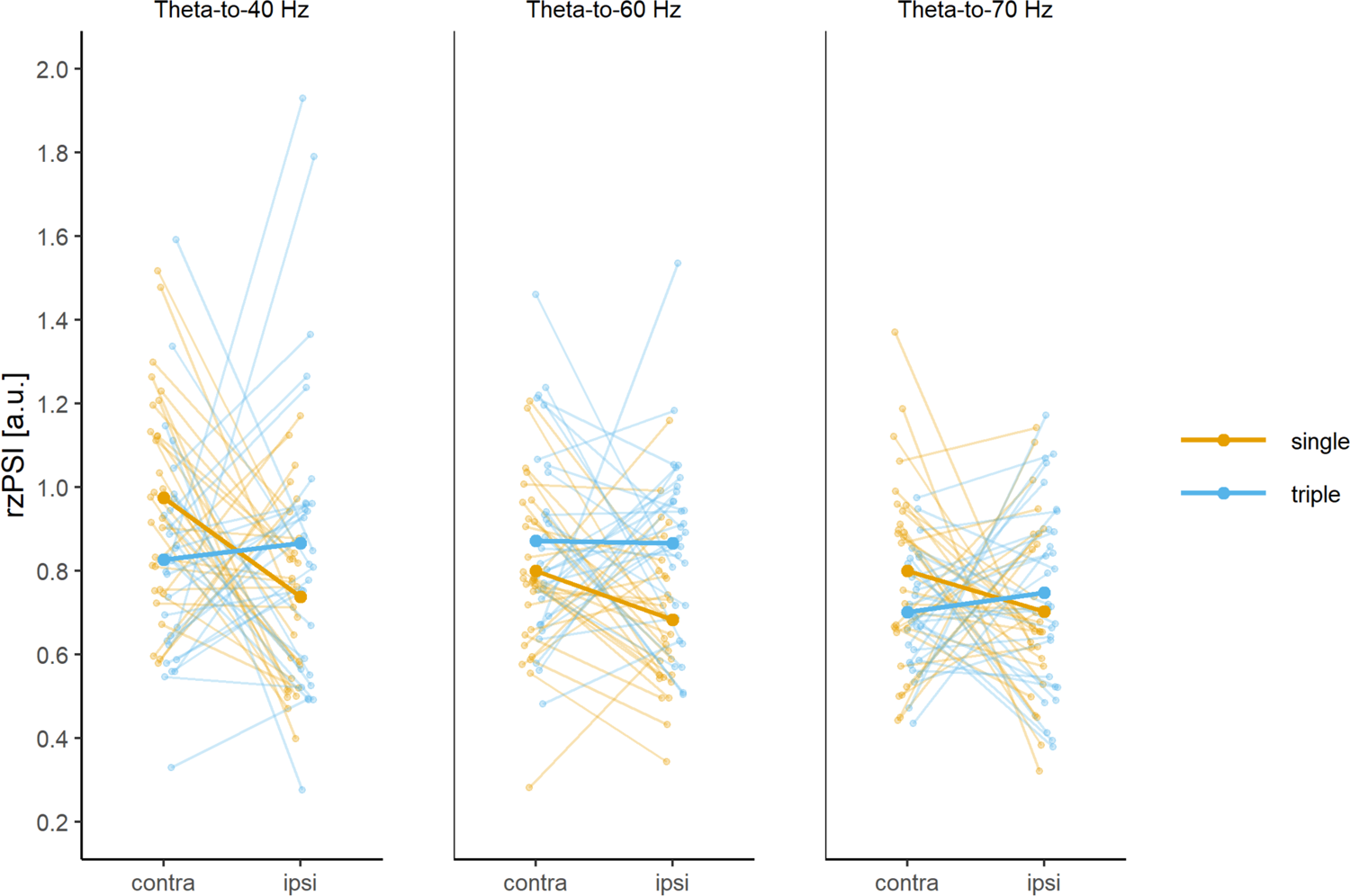
Cross-frequency phase synchronization indices (Rayleigh’s z-transformed; rzPSIs), measuring the consistency of theta-gamma phase difference, from the right hemispheric ROI in the time window 150-200 ms after visual search display onset. RzPSIs are displayed separately for single or triple template conditions (in color), for contralateral or ipsilateral target locations (on the x axis) and for theta-to-40 Hz, theta-to-60 Hz or theta-to-70 Hz cross-frequency synchronization (in separate panels). Single-subject indices (as thin lines) are overlayed by group averages (as thick lines).

For an illustration of the whole time-series for rzPSIs from the right hemispheric ROI, figure 6 shows the descriptives of group average rzPSIs (averaged across theta-to-40 Hz, theta-to-60 Hz or theta-to-70 Hz cross-frequency synchronization) from the right hemispheric ROI for post-stimulus time windows of 50ms length and for a pre-stimulus baseline.

**Figure 6.**
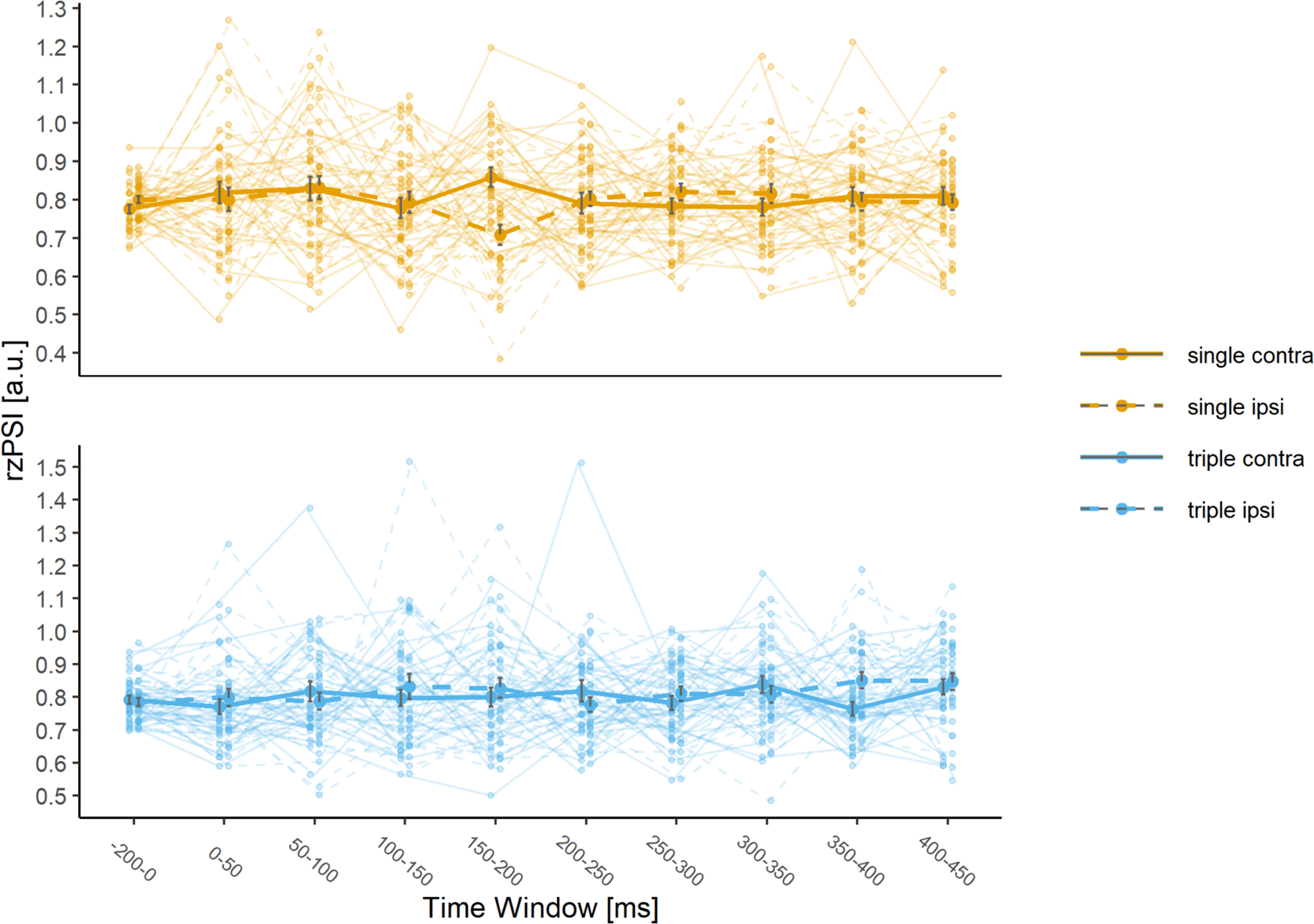
Cross-frequency phase synchronization indices (Rayleigh’s z-transformed; rzPSIs), measuring the consistency of theta-gamma phase difference, from the right hemispheric posterior ROI in windows of 50ms length, starting at stimulus onset 0ms up to 450 ms, and in a 200ms pre-stimulus baseline. Group averaged rzPSIs are shown separately for single or triple template conditions (in color and in separate panels) and for contralateral or ipsilateral target locations (as line-type). Indices are averaged across theta-to-40 Hz, theta-to-60 Hz or theta-to-70 Hz cross-frequency synchronization. Single-subject indices (as thin lines) are overlayed with group averages (as thick lines) and standard errors.

#### Theta-gamma phase synchronization for the left hemispheric ROI

For rzPSIs from the left hemispheric ROI in the time window 150-200 ms after visual search display onset (see supplemental materials S2, figure S2.1), a summary of model fit for the fixed effects COND (single, triple), TARG (contralateral, ipsilateral), CFS (theta-to-40 Hz, theta-to-60 Hz, theta-to-70 Hz), interactions between them, and a single random-effects term for the intercept of the individual subjects can be seen in the supplemental materials S2 in table S2.1.

Unlike the results from the right hemispheric ROI, for rzPSIs from the left hemispheric ROI, no contrast exceeded the threshold of absolute t values larger than 1.96. The grand mean rzPSIs have an estimate of 0.81 as represented by the intercept. Note that the by subject variation beyond the variability induced by the residual error was estimated as zero, i.e. the random effects matrix was singular for this model. However, since dropping a by-subject random effect of zero will have no effect on the fixed effect estimates, it was kept in the model.

For an illustration of the whole time-series for rzPSIs from the left hemispheric ROI, figure S2.2 in the supplemental materials S2 shows the group average rzPSIs (averaged across theta-to-40 Hz, theta-to-60 Hz or theta-to-70 Hz cross-frequency synchronization) from the left hemispheric ROI for post-stimulus time windows of 50ms length and for a pre-stimulus baseline.

### Control analyses

We conducted several control analyses in order to investigate whether the critical interaction between COND and TARG that we found for theta-gamma cross-frequency synchronization in the right-hemispheric ROI is frequency-specific and to exclude a spurious effect relying on evoked responses. These were conducted only for the right-hemispheric ROI because the critical effect was exclusively found for this ROI.

First, to control for the possibility that cross-frequency synchronization is rather between gamma and broadband lower frequencies in general than specifically between gamma and the theta frequency range, the same analyses as in the main analysis were carried out, but for cross-frequency phase synchronization indices between gamma frequencies and the alpha frequency range. If alpha-gamma phase synchronization showed the same pattern of results as theta-gamma cross-frequency synchronization, specifically the critical interaction from the main analysis, the effect would not be frequency specific. Next, to control for the possibility of spurious effects of theta-gamma phase synchronization due to simultaneous but unrelated evoked activity in response to probe presentation in both theta and gamma frequency bands, we analysed spectral amplitudes as well as the phase locking factor (PLF) for theta and gamma frequencies. If a simultaneous increase in spectral amplitudes or a simultaneous phase resetting in response to stimulus onset can be found at both theta and gamma frequencies, this could lead to artificial cross-frequency phase synchronization despite the two frequencies not interacting with each other. This would be the case if spectral amplitudes or rzPLFs for theta and gamma frequencies showed the same pattern of results as the main analysis, specifically the critical interaction. Finally, an analysis using surrogate data was performed. For spurious effects that rely on evoked responses, they should occur at a fixed latency, so surrogate data and real data should show the same pattern of rzPSI estimates, whereas for real effects that are not driven by phase-locking to stimulus onset, real data should show larger rzPSIs estimates than surrogate data.

#### Alpha-gamma phase synchronization for the right hemispheric ROI

To investigate the frequency specificity of the observed interaction between COND and TARG that we found for theta-gamma cross-frequency synchronization in the right-hemisheric ROI in the main analysis, the same analyses were carried out for cross-frequency phase synchronization between gamma frequencies and the alpha frequency range. Thus, the central frequency of interest for this control analysis was at 9.25 Hz (7.40-11.10 Hz) in order to obtain phase estimates from the alpha frequency range. All following analysis steps were identical to the previously described steps for the main analysis of theta-gamma phase synchronization (see methods section for details).

Contrary to the effects observed for the main analysis, in the control analysis for alpha-gamma rzPSIs from the right hemispheric ROI, no contrast exceeded the threshold of absolute t values larger than 1.96. The grand mean rzPSIs have an estimate of 0.8 as represented by the intercept (see supplemental materials S3: figure S3.1 & S3.2 for data visualisation and table S3.1 for a summary of model fit).

#### Amplitudes and PLF for the right hemispheric ROI

To control for the possibility of spurious effects of theta-gamma phase synchronization due to evoked activity in response to probe presentation, we further analysed spectral amplitudes for theta and gamma frequencies. Spectral amplitudes were calculated as the wavelet coefficients’ real values which were then averaged across trials for the same frequencies of interest as in the main analysis. As for the main analysis, we computed averages of amplitudes for time windows of 50ms length and for a pre-stimulus time window of 200 ms. We then also calculated the phase-locking factor (PLF; transformed using Rayleigh’s Z; rzPLF = n*PLF^2) for theta and gamma frequencies separately. This was done to analyse the inter-trial consistency of phase-locking relative to stimulus onset within both frequency bins. For this, their phase values were extracted as in the main analysis (see methods section for details). PLFs were then calculated as the average vector length of these phase values (Bonnefond & Jensen, 2012; Tallon-Baudry et al., 1996), which were also averaged into time windows and transformed using Rayleigh’s Z as in the main analysis.

Thus, for demonstrating that the observed interaction between COND and TARG for theta-gamma cross-frequency synchronization in the right-hemisheric ROI in the main analysis is not an artefact from filtering an evoked response, the same analyses were carried out for theta amplitudes and for gamma amplitudes as well as theta and gamma phase-locking factors from the right hemispheric ROI. While the three gamma bands’ spectral amplitudes as well as their phase-locking factors were calculated separately, they were averaged before entering them into the model because the critical effect in the main analysis did not interact with gamma frequency dependent differences. So for gamma frequencies, both these control analyses were conducted for the average of all three gamma bands.

In the control analysis for theta amplitudes, no contrast exceeded the threshold of absolute t values larger than 1.96. The grand mean theta amplitudes have an estimate of 2.4 (see supplemental materials S6: figure S6.1 & S6.2 for data visualisation and table S4.2 for a summary of model fit). Similarly, in the control analysis for gamma amplitudes, all contrasts remained below the threshold of absolute t values of 1.96. Here, the intercept indicated that the grand mean gamma amplitudes have an estimate of 3.2 (see supplemental materials S7: figure S7.1 & S7.2 for data visualisation and supplemental materials table S7.6 for a summary of model fit). The visualisation of the whole time-series from the theta and gamma band amplitudes from the right hemispheric ROI illustrates that there is no simultaneous increase in both bands in response to stimulus onset. In contrast to theta amplitudes, gamma amplitudes did not show any stimulus-locked increase. Note that while these analyses were conducted for te frequencies of interest, an overview of phase locking values for all Morlet wavelets is presented in figure S4.5 of the supplemental materials S4

In the control analysis for theta rzPLFs, the intercept indicated that the grand mean theta rzPLFs have an estimate of 38. The effect of COND (12, t value = 5.8) indicated that theta rzPLFs are larger in single than triple template conditions (see supplemental materials S5: figure S5.1 & S5.2 for data visualisation and table S5.1 for a summary of model fit). However, in the control analysis for gamma rzPLFs, no contrast exceeded the threshold of absolute t values larger than 1.96. Here, the grand mean gamma rzPLFs have an estimate of 0.95, as indicated by the intercept (see supplemental materials S5: figure S5.3 & S5.4 for data visualisation and table S5.2 for a summary of model fit). When comparing the illustration of the whole time-series from the theta rzPLFs and for gamma rzPLFs from the right hemispheric ROI, it can be seen that gamma rzPLFs did not show a stimulus-locked resetting of phase. Similar to the amplitudes control analysis, there is no indication for a simultaneous increase of phase-locking of both bands in response to stimulus onset.

#### Theta-gamma phase synchronization on surrogate data

For the control analysis using surrogate data, the cross-frequency phase differences were calculated between gamma in a given trial and theta shifted for one trial, resulting in trial-shuffled cross-frequency phase synchronization indices (Rayleigh’s z-transformed; rzPSIs). Importantly, results for surrogate data are not at all similar to those obtained from the analysis for the real data. The critical interaction between COND and TARG from the main analysis (0.18, t = 3.6, see figure 4B) was not reproduced for the analysis on surrogate data (−0.06, t=1.11). And a main effect TARG as in the main analysis (0.062, t = 2.5) was also not present in the analysis on surrogate data (−0.01, t=-0.48), nor was any other effect from the main analysis. The only substantial effects in the model for surrogate data included the contrast CFSTheta60:Theta40: The model showed both a 3-way interaction effect (−0.31, t= −2.55) and a main effect (−0.09, t=2.87) for the CFSTheta60:Theta40 contrast. All other effects were not substantial (t values below 1.96). For comparison with the main analysis on real data, a summary of model fit can be found in supplemental materials S7, including figures for visualization of the surrogate data in the TOI and along the whole time series.

## Discussion

Cross-frequency synchronization between theta and gamma band EEG activity has been proposed to serve matching processes between prediction and sensation in visual perception (Sauseng et al., 2010, 2015). In this study, we investigated how these electrophysiological correlates of memory matching are affected by the number of activated internal templates which can be compared to incoming sensory information. To perform the visual search task of this experiment, one has to hold in mind a template of a single or of multiple targets’ visual properties so that it can be matched with the incoming stimulus. We expected to find stronger transient theta phase to gamma phase synchronization at posterior sites that are contralateral relative to target location compared to ipsilateral targets around 150-200ms after search display presentation in the single template condition, but less so in the triple template condition.

In line with this, we found stronger theta-to-gamma phase synchronization in this early time window at a ROI in the right middle occipital gyrus, elicited by targets presented in the contralateral hemifield than by ipsilateral targets. An increase in theta to gamma phase synchronization contralateral relative to ipsilateral to target locations is well in line with what can be expected due to the lateralized organization of the visual system. This difference between contra- and ipsilateral target locations was larger in the single template condition than in the triple template condition. Specifically, in the single template condition, theta to gamma phase synchronization was higher for contralateral targets than for ipsilateral targets. This was not the case in the triple template condition. Thus, our data lend support to our hypothesis that memory matching was less precise in conditions where one out of three mental templates had to be matched with a target, whereas single template conditions enabled efficient memory matching. Note that these effects were only present in the analysis for the right-hemispheric region of interest, whereas the separate analysis for the left hemispheric ROI did not yield any substantial effects. While we find a clearly right-lateralized effect, there are some previous studies reporting bilateral effects of stronger theta-gamma phase synchronization; e.g. in a cued visual attention task (Sauseng et al., 2008). However, in another study, Holz and colleagues (2010) found a right-lateralized effect of CFS in a visual delayed-match-to-sample task. Similar effects showing a lateralized theta-locked gamma phase synchronization for memory loads 3-4 in visual working memory are also reported by Sauseng and colleagues (2009). Other studies have also found primarily right-hemispheric brain areas to be relevant for visual search. For example, activity enhancements were reported in bilateral superior parietal cortex, but only extending into the intraparietal sulcus of the right hemisphere during visual search compared to overt orienting, (Nobre et al., 2003) and activation was completely right-lateralized for monitoring functions in visual search (Vallesi, 2014). More evidence for clear right-hemispheric dominance for search organization comes from lesion-studies, for example that lesions in the right parietal, temporal and occipital cortex were related to disorganized search (Ten Brink et al., 2016). This may explain why we only find a larger difference between contra- and ipsilateral target locations in the single template condition than in the triple template condition for right posterior, but not for left posterior regions of interest. While our right-hemispheric results reproduce previous evidence (Holz et al., 2010; Sauseng et al., 2008), the left-hemispheric results from these studies appear to be more variable overall: In the current data, we basically find no effect, whereas previously, both a left-hemispheric reversal of the effect in a visual delayed-match-to-sample task (Holz et al., 2010), as well as a similar effect in the same direction as in right hemisphere in a cued visual attention task (Sauseng et al., 2008) have been reported. Taken together, the left-hemispheric patterns of results seem to depend more on the specific task paradigm at hand. We will therefore focus on discussing the right-hemispheric results in the following.

This evidence from the single template condition in the current visual search paradigm corroborates previous findings showing a higher transient phase synchronization between posterior theta and gamma activity in cases where our expectancies match the actual visual input than in case of a non-match (Holz et al., 2010; Sauseng et al., 2008). Thus, our data from the single template condition fit well into the proposed framework that could well account for the activation of mental templates from working memory and their comparison with sensory input (Sauseng et al., 2010, 2015), which proposes that cross-frequency phase synchronization between theta and gamma frequencies early after target presentation can be regarded as a candidate neural mechanism underlying the matching of mental templates from working memory with sensory input. Here, the typically observed increase of fronto-parietal phase-coupling in the theta band during anticipation of a specified visual target is suggested to reflect the active presentation of a mental template in working memory, controlled by frontal resources and replayed into higher visual areas. Then subsequently, a posterior phase resetting of theta band oscillations is assumed to enable the transient cross-frequency coupling synchronization with high frequency activity in the gamma band range repeatedly found in a time window around 150 ms after target presentation. While potential alternative explanations will be discussed later, converging evidence supporting this view exists, suggesting that frontal low-frequency oscillations are indeed crucially involved in the top-down control during working memory tasks through coherence and cross-frequency interaction in fronto-parietal networks (for recent reviews, see de Vries et al., 2020; Karakaş, 2020; Klink et al., 2020; Palva & Palva, 2018).

Conversely, the triple template condition does not reveal such dynamics early after target presentation. A plausible interpretation of this results is that in the triple template search condition multiple templates are held sequentially in working memory; and that in a given trial depending on whether the sequence’s first, second, or third mental template could be matched to the current visual input, memory matching occurred relatively early, a bit later or even much later, leading to overall more temporal variability across trials. This is then reflected in the cross-frequency phase synchronization mechanism investigated here. Based on proposals for a limited capacity of human visual working memory, supposedly around three to four items (Luck & Vogel, 2013), one might expect that likewise, we could maintain up to three or four simultaneous search templates for visual search as well. However, there is evidence that not all working memory items influence the guidance of selective attention, but that only active memory items function as an attentional template and directly affect perception, whereas accessory memory items have relatively little influence on visual selection (Olivers et al., 2011). The conclusion here is that working memory items generally compete for the status of ‘attentional template’, which can only be achieved by one item at a time. So, although working memory could store multiple objects, observers could only actively look for one at a time. Thus, multiple-template search should require switching between mental templates or sequentially looking for them. Interestingly, though, Beck and colleagues (2012) propose that observers can concurrently keep two templates active in simultaneous search because when they explicitly asked participants to simultaneously search for two templates, their gaze frequently switched between them without switch costs. Similarly, Hollingworth and Beck (2016) found that even when multiple templates were kept in mind, a distractor in a visual-search task captured attention more when it matched the template(s), and proposed that multiple templates can guide attention simultaneously; but see also van Moorselaar et al. (2014) or Frătescu et al. (2019) where in contrast, such memory-driven capture was reported only for single templates, demonstrating that this does not hold in all situations. However, evidence showing clear switch costs for selection has been reported when only one out of two potential targets was available, suggesting that observers cannot actively search for multiple objects if they are not able to freely choose the target category (Ort et al., 2017, 2018). This is again well in line with the idea that only one search template at a time has priority and will guide visual attention (Olivers et al., 2011) and Ort and colleagues argue that both lines of evidence can be explained from a reactive versus proactive cognitive control framework (Braver, 2012). In this framework, when multiple targets are all available for search, participants can proactively prepare for any target, resulting in a lack of switch costs. Conversely, when only one of multiple possible targets is present for search, the currently displayed target might not match the target that the participant anticipated. Reactive control would follow this conflict, leading to increased processing times.

This latter case is similar to the current study’s triple template condition, where only one of three possible targets was presented for search. While the design of our task does not allow for the analysis of inter-trial switch costs, we do see significantly fewer correct responses as well as behavioural slowing in response times to targets in the triple template blocks compared to targets in single template blocks. Slower and less accurate search performance has also been reported during simultaneous search for two targets compared to search for either target alone, indicating that subjects can probably not perform two simultaneous matching processes (Huang & Pashler, 2007; Menneer et al., 2007). So most likely, in a given trial in the current study’s triple template blocks, a currently displayed target possibly did not match the target that the participant anticipated. This means that across trials, the match between memory templates and visual input would occur at varying points in time, which should also be reflected in the neural correlates of memory matching, predicting low estimates of cross-frequency phase synchrony. Conversely, in the single template condition, certainty about the mental template that has to be matched with sensory input was high in each trial and enabled a temporally precise matching process across trials, which predicts higher estimates of cross-frequency phase synchrony. Our data support this interpretation well, showing that theta to gamma phase synchronization, the proposed underlying mechanism of memory matching, was higher for contralateral targets than for ipsilateral targets in the single template condition, whereas this was not the case in the triple template condition. Note that as a measure for theta-gamma cross-frequency phase synchrony, we analysed the consistency of phase difference between the two frequencies over trials (Holz et al., 2010; Palva et al., 2005; Sauseng et al., 2008). This measure does not require the two frequencies to be coupled continuously. High estimates of phase synchrony will be achieved when there is a fixed relation between low and high frequencies across trials, independent of absolute phase difference between them and of phase-locking to stimulus of either of them, whereas low estimates will be achieved when phase relations vary over trials.

To discuss the pattern of results in the triple template condition, one might want to speculate about how the memory matching mechanism investigated here might rely on pre-stimulus working memory retention mechanisms. Generally, cross-frequency interactions between gamma band activity and slower brain waves have frequently been suggested to be involved in multi-item working memory, such as for multi-item working memory retention. A prominent computational model assumes that separate memory items are represented by single gamma waves which are nested into a theta wave (Jensen & Lisman, 1998; Lisman & Idiart, 1995). It is hypothesized that with this mechanism, multiple memory items (gamma waves nested into a theta cycle) can be actively held in parallel in working memory. In the light of this framework, one would assume that before search display presentation, in the triple template condition, the three mental templates would each be represented by separate gamma cycles nested into a theta wave one after another. Thus, upon search display presentation, it would be crucial as to whether the first, second or third item (gamma wave) incidentally matches with the one on the search display, leading to a temporal variability in the range of two gamma cycles. Another theoretical framework which entails cross-frequency synchronization between theta and gamma as the neural basis for multi-item WM retention argues that during retention, each item is coded by an entire gamma burst, i.e. multiple cycles, nested into a theta wave (Herman et al., 2013; Van Vugt et al., 2014). Following these ideas, there would be a temporal variability of memory matching in the triple template condition in the range of two theta cycles, depending on whether visual input matches with the first, second or third item (gamma burst). In this study, grand mean reaction time differences in the triple template condition were longer than reaction times in the single template condition by about 190 ms, which is in the range of a theta cycle. Speculatively, this would fit rather well with the predictions derived from the latter framework, where due to the expected temporal variability of two theta cycles, average reaction times would be expected to be around the length of one theta cycle longer when one out of three potential targets can be matched with visual input. Note, however that our task instructions had not emphasized a speeded but rather an accurate response, so we cannot draw strong conclusions here. To examine these predictions more closely, studies with a more precise measurement of response times would be required.

Although a sequential matching process seems to be a rather plausible interpretation of the observed low estimates of cross-frequency phase synchrony in the triple template condition, there may be alternative explanations to this. For example, a similar pattern of results could be obtained when cross-frequency phase-relations exhibit overall more temporal variability across trials due to differences in the source of EEG activity when several templates have to be processed. Or, for example, low phase synchronisation estimates would be expected if memory matching in the triple template conditions happened with great temporal variability and if it happened consistently later than in the single template conditions. Also, an unspecific difference between the conditions, such as larger neural noise could have resulted in low estimates in the triple condition. We cannot rule out these possibilities. Alternatively assuming a parallel mode, would predict that multiple templates interacted in parallel with sensory input; however, this would come at costs due to mutual competition, leading to a delay in target selection (Ort et al., 2019; Ort & Olivers, 2020). For the matching phase, this may mean that when one out of multiple targets must be found, the matching process would happen later than for a single possible target, but consistently, with low temporal variability across trials. In this case, we would have expected to observe a slightly later effect, but with high estimates of phase synchrony similar to those in the single template condition. Descriptively, we find no indication for something like this in our data. Finally, it could be that participants might have had less precise, low fidelity templates in the triple template condition. One possibility is that template fidelity in multi-item retention could influence theta phase. We recently argued that an increased memory fidelity could be an explanation for the empirically observed increased memory capacity by slowing down theta waves (Vosskuhl et al., 2015; Wolinski et al., 2018): In the light of theoretical models proposing that single visual items are neuronally represented by entire gamma bursts nested into theta waves for multi-item retention (Herman et al., 2013; Van Vugt et al., 2014), an increased memory fidelity could be an explanation for these findings because longer gamma bursts, representing a template with more fidelity, outweigh the slower rate of memory re-activation (Sauseng et al., 2019). Thus, a “let’s see if something looks familiar” search mode due to overall lower template fidelity would predict shorter theta cycles than a mode where there is an active top-down set for the target properties. This would predict that theta cycles may have been shorter in the triple, compared to the single template condition. But even if theta frequency was sped up, this should not automatically lead to attenuated theta-gamma phase synchronization. Based on the nature of this measure, only increased temporal jitter should lead to that effect. Therefore, our pattern of results does not support the idea of fidelity differences of the conditions. However, another possibility is that familiarity matching processes could have an entirely different neural signature than template matching processes. Since this should not be reflected in the investigated cross-frequency coupling index, we cannot exclude this possibility.

Yet, we assumed that visual search for the kind of complex targets we used in this study would have relied on an active attentional template in working memory (Gunseli et al., 2014). The abstract symbols that we used as target and distractor stimuli were rather complex and all unknown to the observers. However, since it was not possible to ask participants to memorize a trial-by-trial changing target and, thus, a training beforehand was necessary, it is likely that participants may have stored those memorized target(s) in long-term memory before the start of our task. Given that the task was still relatively difficult (task performance was on average 86% and 63% correct responses in the single and triple condition, respectively), however, we think that rather than being stored passively in long-term memory, it is more likely that the template had to be activated in working memory for successful task performance (Ruchkin et al., 2003). In an ERP study, Gunseli and colleagues (2014) reported evidence for a larger LPC component when an effortful, as opposed to an efficient search is anticipated, indicating that participants tried maintaining an attentional template in working memory with greater effort. In other words, this suggests that for effortful search an increased working memory effort for maintaining the template in working memory may be required. Based on our participant’s feedback and task performance, it seems plausible that our task was experienced as quite challenging; and the strategies that were reported in the personal feedback indicate that they tried maintaining a target template vividly. However, we cannot claim that our task was a pure working memory paradigm, as clear long-term memory involvement exists. Thus, in order to strengthen the argument for matching of templates from working memory, the paradigm could be adjusted to a trial-by-trial target cueing, without prior training in the future. Yet, EEG studies suggest that even in design where targets are not trained beforehand, but changed on a trial-by-trial basis, the attentional template is learned after repeated search for the same target, as evidence for decreased CDA and LPC components with target repetition was found (Carlisle et al., 2011; Gunseli et al., 2014). Based on this, it is proposed that an attentional template which is initially stored in working memory can be transferred to long-term memory when the target is repeated. Additionally, contextual cueing effects in visual search (e.g. Zinchenko et al., 2020) can be explained through storing spatial target-distractor relations as templates in long-term memory after they have been repeatedly encountered. This would mean that even in a search paradigm where targets are cued in each trial, the involvement of long-term memory cannot be entirely excluded.

While the evidence and framework we build on has its focus on how mental templates interact with visual information before and until the match between stimulus-related information and memory contents happens (Sauseng et al., 2010, 2015), memory matching is probably one of several steps that are assumed to take place within the selection stage of visual search (for review, see Eimer (2014) and Ort & Olivers, (2020). For example, the ‘match-and-utilization model’ focuses both on the step of the match between stimulus-related information and memory contents as well as the step of utilization, where the result of this match or mismatch is then ‘read out’, which could then result in the updating of memory, the selection of behavioural responses and the reallocating of attention (Herrmann et al., 2010). Or, from the point of view of predictive coding theories which assume that top-down predictions are matched to incoming sensory inputs across different levels of the cortical hierarchy, it is assumed that a prediction-error signal is fed forward along the cortical hierarchy and used to update top-down predictions (Friston, 2005). For the template-matching visual input to win the competitive race over other visual input, an increase in attention towards the identity or spatial location of memory-matching visual inputs is quite likely. We cannot exclude that such mechanisms are contributing to the observed effects in our study.

The right-hemispheric effects from our data seem to be specific for theta-gamma phase synchronization since a control analysis for cross-frequency phase synchronization between alpha and gamma did not show similar results. Yet, for alpha-gamma phase synchronization there was an interaction between template condition and target location for the left hemispheric posterior source. However, the direction of this effect (stronger ipsilateral PSI) was contrary to what would be expected due to the lateralized organization of the visual system.

Previous studies have reported similar cross-frequency interaction either between theta and gamma frequencies around 30-50Hz in a cued visual attention task (Sauseng et al., 2008) or between theta and higher gamma activity (Holz et al., 2010; Sauseng et al., 2009). In the current data from our visual search paradigm, it seems that the difference between right posterior rzPSIs for Theta-to-70Hz and for Theta-to-60Hz is smaller for single than triple template conditions; whereas the difference for Theta-to-60Hz and for Theta-to-40Hz seems to be larger for single than triple template conditions. However, the critical effect between template condition and target location in the main analysis did not interact with gamma frequency dependent differences. This speaks rather for a broadband gamma effect than a selective effect of theta and a narrow gamma sub-band in the current visual search paradigm. The critical interaction from the main analysis seems to be rather frequency-specific to theta-gamma phase synchronization, however, because contrary to the effects observed for the main analysis, all contrasts remained below threshold in a control analysis with alpha-gamma phase synchronization.

To ensure that differences between conditions were not based on merely different trial counts in the single and triple template condition, PSI values were transformed using Rayleigh’s z transform (Cohen, 2014). Naturally, this does not eliminate the difference in signal-to-noise ratio between conditions, which was most likely lower in the triple template condition. Note, however, that this pattern of results was found even though in both conditions, only trials with a correct response for which we assume that successful memory matching must have taken place at some point were used to calculate theta-gamma phase synchronization indices. Additionally, we found that in a control analysis where we drew a random subset of the same number of trials in the condition with fewer trials before calculating rzPSIS on these trial-matched data, results were very similar to those obtained from the main analysis based on all trials. Importantly, the critical interaction between COND and TARG from the main analysis was reproduced and showed the same pattern of results, namely that the single template condition showed larger estimates for contra compared to ipsilateral targets, whereas this was not the case for the triple condition.

Because spurious effects of theta-gamma phase synchronization might arise due to evoked activity in response to probe presentation, we analysed amplitudes and the phase locking factor for theta and gamma frequencies to control for this. If both frequencies showed a simultaneous increase in amplitudes or a simultaneous phase resetting in response to stimulus onset, the data might indicate artificial cross-frequency phase synchronization in the absence of true interactions between the two frequencies. However, none of these control analyses showed a simultaneous increase in amplitudes or a simultaneous phase resetting in response to stimulus onset nor a similar pattern of results as the main analysis. Thus, the results from these control analyses rather indicate it being implausible that the observed interaction between COND and TARG for theta-gamma cross-frequency synchronization in the right-hemisheric ROI in the main analysis, is due to an artefact from simultaneous evoked activity in response to probe presentation in both theta and gamma frequency bands. This is also confirmed by a control analysis on surrogate data, demonstrating that trial-shuffled cross-frequency phase synchronization indices did not show similar results to the real data, which should have been the case if the observed effect of theta-gamma phase synchronization 150–200ms after probe presentation in the real data was generated through an evoked response.

## Conclusion

Taken together, our data lend support to the hypothesis that neuronal networks operating at theta and gamma frequency do become more synchronized in phase during an early time window following visual search display onset, when a single template has to be retained compared to triple template conditions. This adds to previous theoretical accounts that have strongly argued for a transient synchronization between theta and gamma phase over posterior electrode sites as a neural correlate of matching of incoming sensory information with memory contents from working memory (Sauseng et al., 2010, 2015). We interpret this as showing that while a single mental template enables precise memory matching, limitations in this matching process occur during multiple template search. These could be explained by sequential attentional templates (Lisman & Idiart, 1995; Olivers et al., 2011; Van Vugt et al., 2014), however, other task paradigms combining multiple template search with the investigation of target switch costs ought to corroborate this. For future studies, it would be interesting to investigate the temporal dynamics of such matching processes during the acquisition and consolidation phase of attentional templates. Studying more naturalistic contexts of template to input matching where, for example, templates are acquired via learning, could further illuminate the involvement of cross frequency interactions in template to input matching.

## Author Contributions

**Anna Lena Biel:** Conceptualization, Methodology, Software, Formal analysis, Investigation, Data Curation, Writing – Original Draft, Visualization, Project administration; **Tamas Minarik:** Conceptualization, Writing – Review & Editing; **Paul Sauseng:** Conceptualization, Methodology, Validation, Resources, Writing – Review & Editing, Supervision, Funding acquisition

## Acknowledgements

This work was funded by Deutsche Forschungsgemeinschaft (DFG) SA 1872/2-1. We thank Eva Margraf and Patricia Christian for their support with data collection and Marleen Haupt for valuable discussions.

