## supplemental materials for "EEG Cross-Frequency Phase Synchronization as an Index of Memory Matching in Visual Search"

### S1 Theta-gamma phase synchronization at right hemispheric ROI

**Table S1.1.** Summary of model fit for theta-gamma phase synchronization indices (rzPSIs) from the right hemispheric ROI in the time window 150-200 ms after visual search display onset.

Linear mixed model fit by REML

REML criterion at convergence: 19

Scaled residuals:

| Min | 1Q | Median | 3Q | Max |
|---|---|---|---|---|
| -2.55 | -0.69 | -0.09 | 0.59 | 4.41 |

Random effects:

| Groups | Term | Std.Dev. |
|---|---|---|
| SUBJ | (Intercept) | 0.03310 |
| Residual | | 0.23236 |

Number of obs: 348, groups: SUBJ, 29.

Fixed effects:

| | Estimate | Std. Error | t value |
|---|---|---|---|
| (Intercept) | 0.8 | 0.014 | 57 |
| CONDSinglevTriple | -0.03 | 0.025 | -1.2 |
| TARGContravIpsi | 0.062 | 0.025 | 2.5 |
| CFSTheta60vTheta40 | -0.046 | 0.031 | -1.5 |
| CFSTheta70vTheta60 | -0.068 | 0.031 | -2.2 |
| CONDSinglevTriple:TARGContravIpsi | 0.18 | 0.05 | 3.6 |
| CONDSinglevTriple:CFSTheta60vTheta40 | -0.14 | 0.061 | -2.3 |
| CONDSinglevTriple:CFSTheta70vTheta60 | 0.15 | 0.061 | 2.5 |
| TARGContravIpsi:CFSTheta60vTheta40 | -0.038 | 0.061 | -0.62 |
| TARGContravIpsi:CFSTheta70vTheta60 | -0.035 | 0.061 | -0.58 |
| CONDSinglevTriple:TARGContravIpsi:CFSTheta60vTheta40 | -0.17 | 0.12 | -1.4 |
| CONDSinglevTriple:TARGContravIpsi:CFSTheta70vTheta60 | 0.034 | 0.12 | 0.28 |

*Note.* The model includes a random-effects term for the intercept of individual subjects and the fixed effects COND (Single, Triple), TARG (Contralateral, Ipsilateral), CFS (Theta-to-40 Hz, Theta-to-60 Hz, Theta-to-70 Hz), and interactions between them. Contrast coefficients with absolute *t* values larger than 1.96 can be treated as approximating the 5% significance level. rzPSI = cross-frequency phase synchronization index, Rayleigh's z-transformed.

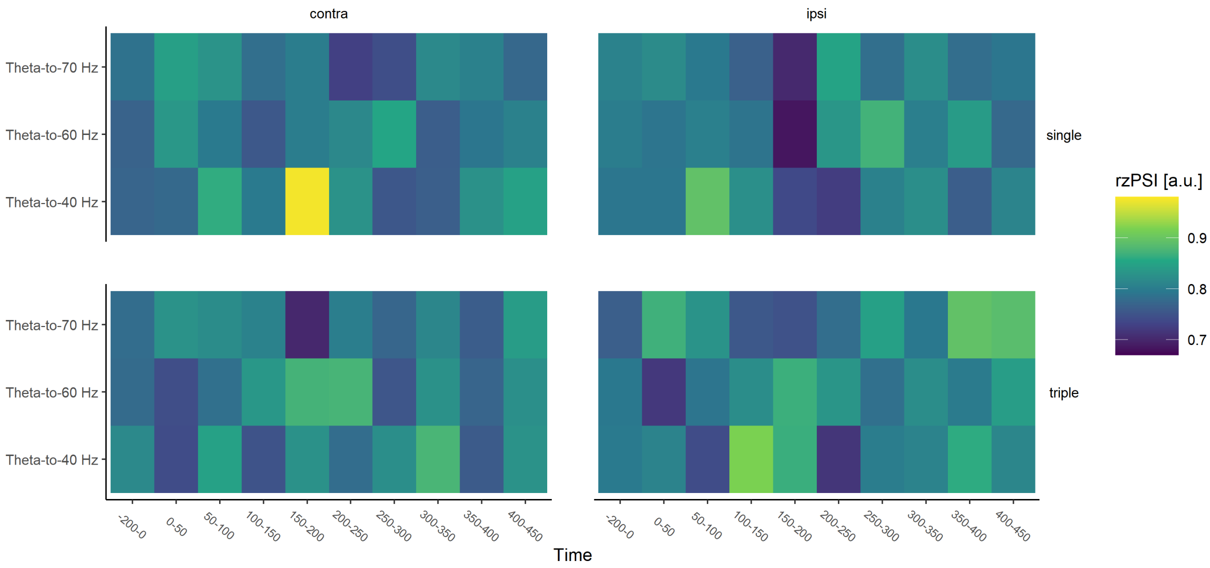

**Figure S1.1.** Cross-frequency phase synchronization indices (Rayleigh's z-transformed; rzPSIs), measuring the consistency of theta-gamma phase difference from the right hemispheric posterior ROI in windows of 50ms length, starting at stimulus onset 0ms up to 450 ms, and in a 200ms pre-stimulus baseline, for theta-to-40 Hz, theta-to-60 Hz and theta-to-70 Hz phase synchronization. Group averaged rzPSIs are shown separately for single or triple template conditions and for contralateral or ipsilateral target locations.

### S2 Theta-gamma phase synchronization at left hemispheric ROI

**Table S2.1.** Summary of model fit for theta-gamma phase synchronization indices (rzPSIs) from the left hemispheric ROI in the time window 150-200 ms after visual search display onset.

Linear mixed model fit by REML

REML criterion at convergence: -14

Scaled residuals:

| Min | 1Q | Median | 3Q | Max |
|---|---|---|---|---|
| -1.97 | -0.72 | -0.11 | 0.65 | 5.04 |

Random effects:

| Groups | Term | Std.Dev. |
|---|---|---|
| SUBJ | (Intercept) | 0.00000 |
| Residual | | 0.22309 |

Number of obs: 348, groups: SUBJ, 29.

Fixed effects:

| | Estimate | Std. Error | t value |
|---|---|---|---|
| (Intercept) | 0.81 | 0.012 | 68 |
| CONDSinglevTriple | -0.0098 | 0.024 | -0.41 |
| TARGContravIpsi | 0.016 | 0.024 | 0.66 |
| CFSTheta60vTheta40 | 0.017 | 0.029 | 0.57 |
| CFSTheta70vTheta60 | 0.0052 | 0.029 | 0.18 |
| CONDSinglevTriple:TARGContravIpsi | -0.043 | 0.048 | -0.9 |
| CONDSinglevTriple:CFSTheta60vTheta40 | -0.019 | 0.059 | -0.33 |
| CONDSinglevTriple:CFSTheta70vTheta60 | -0.013 | 0.059 | -0.22 |
| TARGContravIpsi:CFSTheta60vTheta40 | 0.075 | 0.059 | 1.3 |
| TARGContravIpsi:CFSTheta70vTheta60 | -0.039 | 0.059 | -0.67 |
| CONDSinglevTriple:TARGContravIpsi:CFSTheta60vTheta40 | -0.12 | 0.12 | -1 |
| CONDSinglevTriple:TARGContravIpsi:CFSTheta70vTheta60 | -0.081 | 0.12 | -0.69 |

*Note.* The model includes a random-effects term for the intercept of individual subjects and the fixed effects COND (Single, Triple), TARG (Contralateral, Ipsilateral), CFS (Theta-to-40 Hz, Theta-to-60 Hz, Theta-to-70 Hz), and interactions between them. Contrast coefficients with absolute *t* values larger than 1.96 can be treated as approximating the 5% significance level. rzPSI = cross-frequency phase synchronization index, Rayleigh's z-transformed.

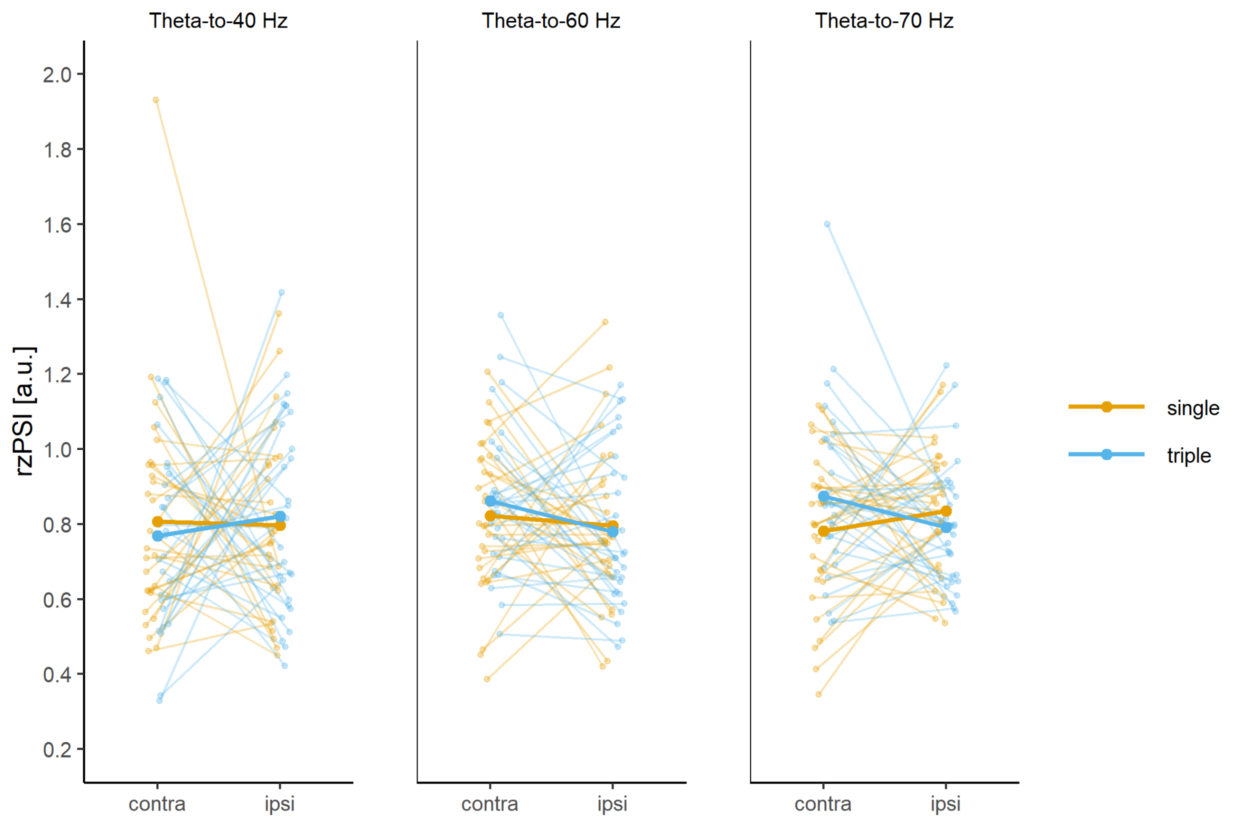

**Figure S2.1.** Cross-frequency phase synchronization indices (Rayleigh's z-transformed; rzPSIs), measuring the consistency of theta-gamma phase difference, from the left hemispheric ROI in the time window 150-200 ms after visual search display onset. RzPSIs are displayed separately for single or triple template conditions (in colour), for contralateral or ipsilateral target locations (on the x axis) and for theta-to-40 Hz, theta-to-60 Hz or theta-to-70 Hz cross-frequency synchronization (in separate panels). Single-subject indices (as thin lines) are overlayed with group averages (as thick lines).

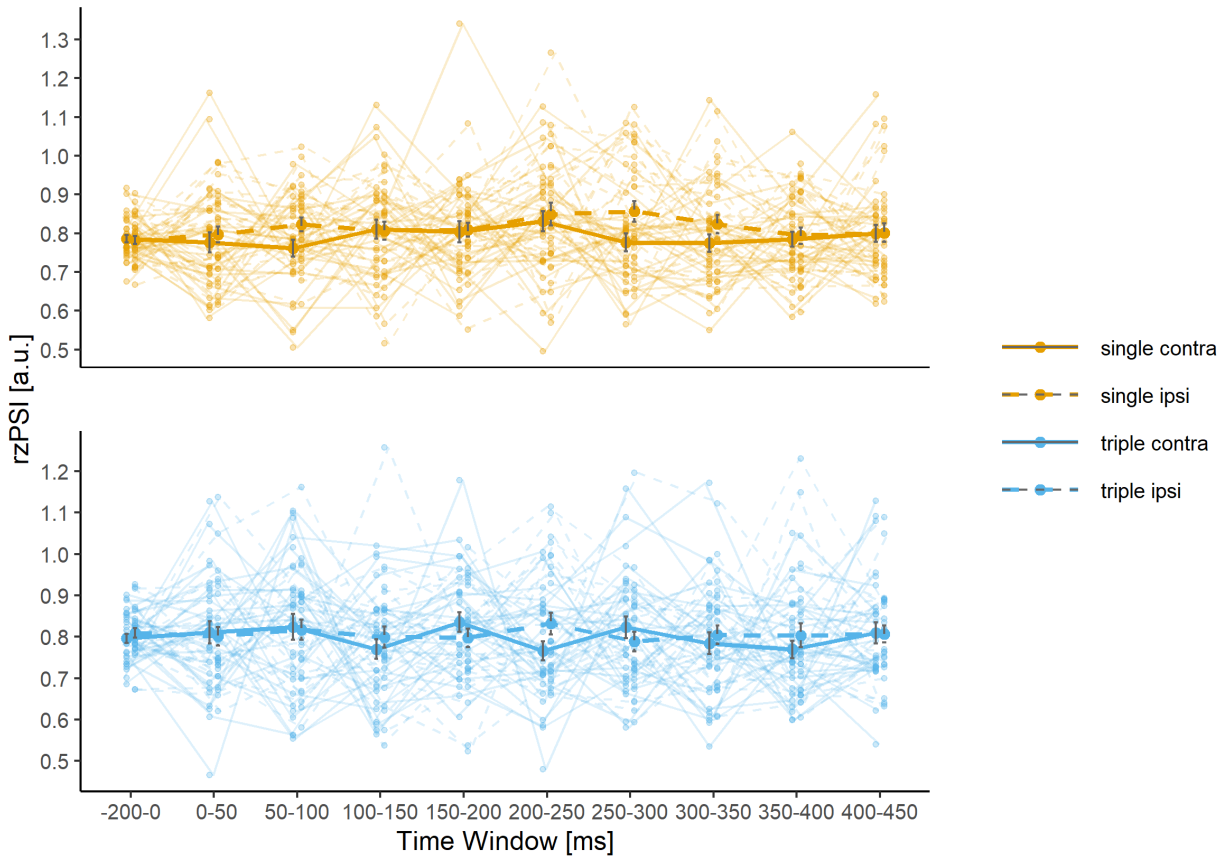

**Figure S2.2.** Cross-frequency phase synchronization indices (Rayleigh's z-transformed; rzPSIs), measuring the consistency of theta-gamma phase difference, from the left hemispheric posterior ROI in windows of 50ms length, starting at stimulus onset 0ms up to 450 ms, and in a 200ms pre-stimulus baseline. Group averaged rzPSIs are shown separately for single or triple template conditions (in color and in separate panels) and for contralateral or ipsilateral target locations (as line-type). Indices are averaged across theta-to-40 Hz, theta-to-60 Hz or theta-to-70 Hz cross-frequency synchronization. Single-subject indices (as thin lines) are overlayed with group averages (as thick lines) and standard errors.

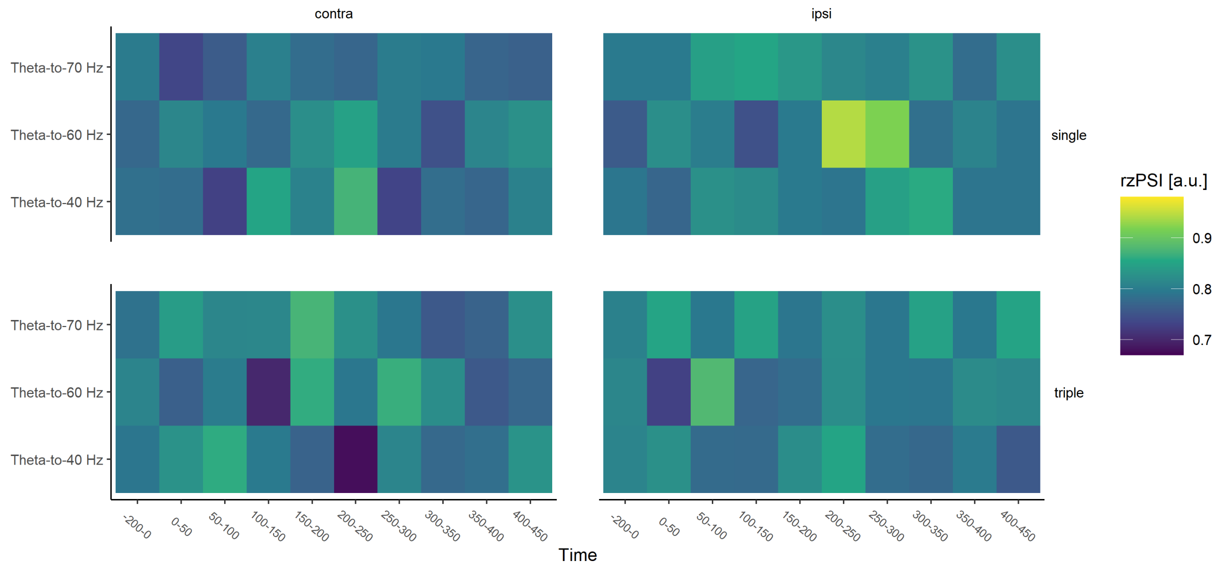

**Figure S2.3.** Cross-frequency phase synchronization indices (Rayleigh's z-transformed; rzPSIs), measuring the consistency of theta-gamma phase difference from the left hemispheric posterior ROI in windows of 50ms length, starting at stimulus onset 0ms up to 450 ms, and in a 200ms pre-stimulus baseline, for theta-to-40 Hz, theta-to-60 Hz and theta-to-70 Hz phase synchronization. Group averaged rzPSIs are shown separately for single or triple template conditions and for contralateral or ipsilateral target locations.

### S3 Control analysis: Alpha-gamma phase synchronization

**Table S3.1.** Summary of model fit for alpha-gamma phase synchronization indices (rzPSIs) from the right hemispheric ROI in the time window 150-200 ms after visual search display onset.

Linear mixed model fit by REML

REML criterion at convergence: -95

Scaled residuals:

| Min | 1Q | Median | 3Q | Max |
|---|---|---|---|---|
| -2.33 | -0.61 | -0.06 | 0.57 | 5.73 |

Random effects:

| Groups | Term | Std.Dev. |
|---|---|---|
| SUBJ | (Intercept) | 0.025446 |
| Residual | | 0.196432 |

Number of obs: 348, groups: SUBJ, 29.

Fixed effects:

| | Estimate | Std. Error | t value |
|---|---|---|---|
| (Intercept) | 0.8 | 0.012 | 69 |
| CONDSinglevTriple | -0.012 | 0.021 | -0.58 |
| TARGContravIpsi | -0.022 | 0.021 | -1 |
| CFSAlpha60vAlpha40 | -0.046 | 0.026 | -1.8 |
| CFSAlpha70vAlpha60 | 0.042 | 0.026 | 1.6 |
| CONDSinglevTriple:TARGContravIpsi | -0.065 | 0.042 | -1.5 |
| CONDSinglevTriple:CFSAlpha60vAlpha40 | 0.003 | 0.052 | 0.059 |
| CONDSinglevTriple:CFSAlpha70vAlpha60 | -0.023 | 0.052 | -0.44 |
| TARGContravIpsi:CFSAlpha60vAlpha40 | 0.048 | 0.052 | 0.94 |
| TARGContravIpsi:CFSAlpha70vAlpha60 | -0.051 | 0.052 | -0.99 |
| CONDSinglevTriple:TARGContravIpsi:CFSAlpha60vAlpha40 | -0.089 | 0.1 | -0.86 |
| CONDSinglevTriple:TARGContravIpsi:CFSAlpha70vAlpha60 | 0.099 | 0.1 | 0.96 |

*Note.* The model includes a random-effects term for the intercept of individual subjects and the fixed effects COND (Single, Triple), TARG (Contralateral, Ipsilateral), CFS (Alpha-to-40 Hz, Alpha-to-60 Hz, Alpha-to-70 Hz), and interactions between them. Contrast coefficients with absolute $t$ values larger than 1.96 can be treated as approximating the 5% significance level. rzPSI = cross-frequency phase synchronization index, Rayleigh's z-transformed.

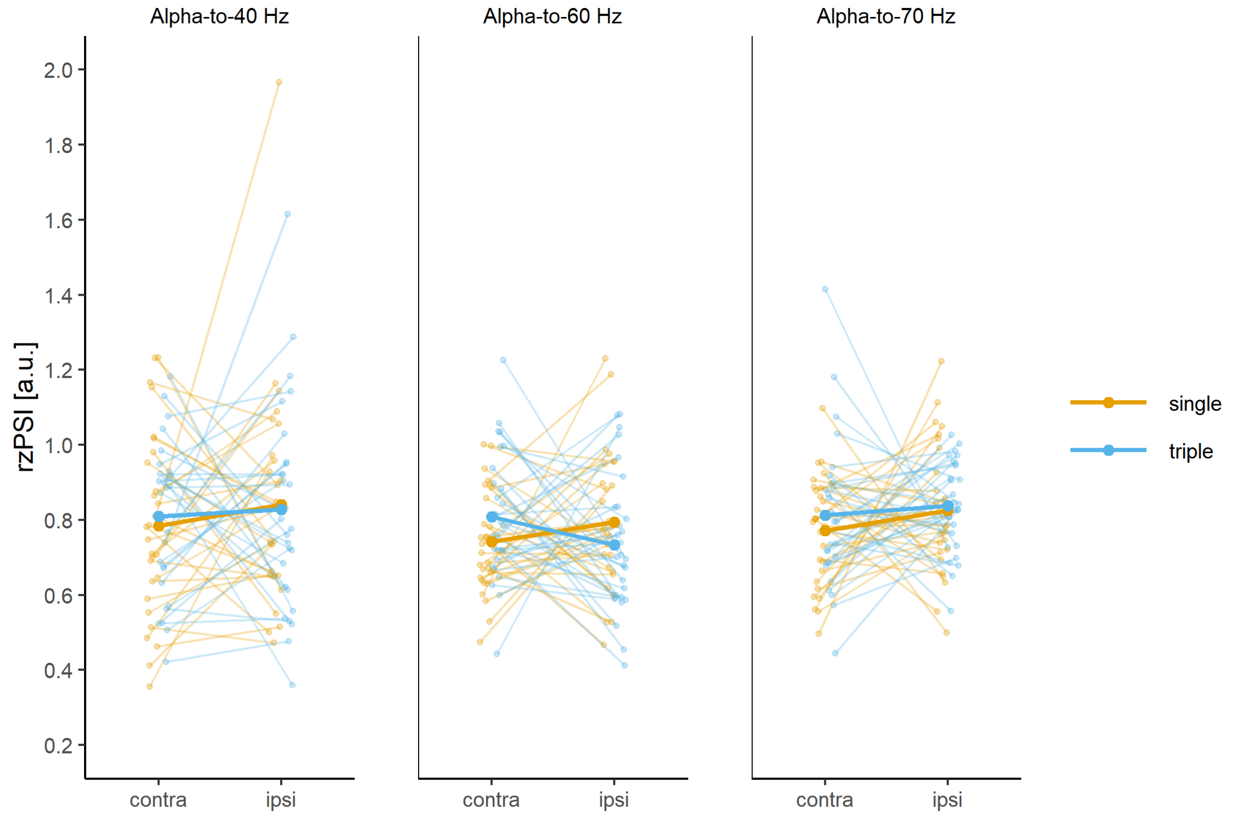

**Figure S3.1.** Cross-frequency phase synchronization indices (Rayleigh's z-transformed; rzPSIs), measuring the consistency of alpha-gamma phase difference, from the right hemispheric ROI in the time window 150-200 ms after visual search display onset. RzPSIs are displayed separately for single or triple template conditions (in colour), for contralateral or ipsilateral target locations (on the x axis) and for alpha-to-40 Hz, alpha-to-60 Hz or alpha-to-70 Hz cross-frequency synchronization (in separate panels). Single-subject indices (as thin lines) are overlayed with group averages (as thick lines).

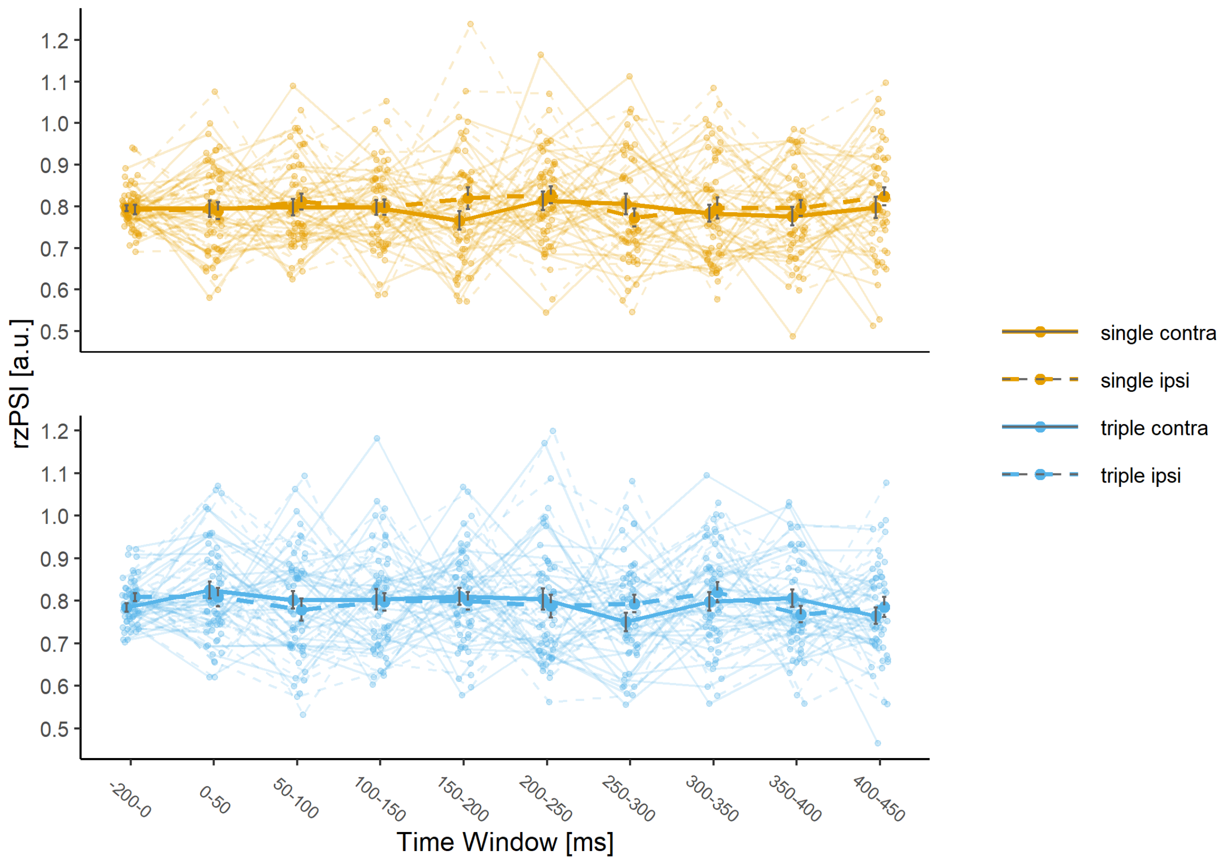

**Figure S3.2.** Cross-frequency phase synchronization indices (Rayleigh's z-transformed; rzPSIs), measuring the consistency of alpha-gamma phase difference, from the right hemispheric posterior ROI in windows of 50ms length, starting at stimulus onset 0ms up to 450 ms, and in a 200ms pre-stimulus baseline. Group averaged rzPSIs are shown separately for single or triple template conditions (in color and in separate panels) and for contralateral or ipsilateral target locations (as line-type). Indices are averaged across alpha-to-40 Hz, alpha-to-60 Hz or alpha-to-70 Hz cross-frequency synchronization. Single-subject indices (as thin lines) are overlayed with group averages (as thick lines) and standard errors.

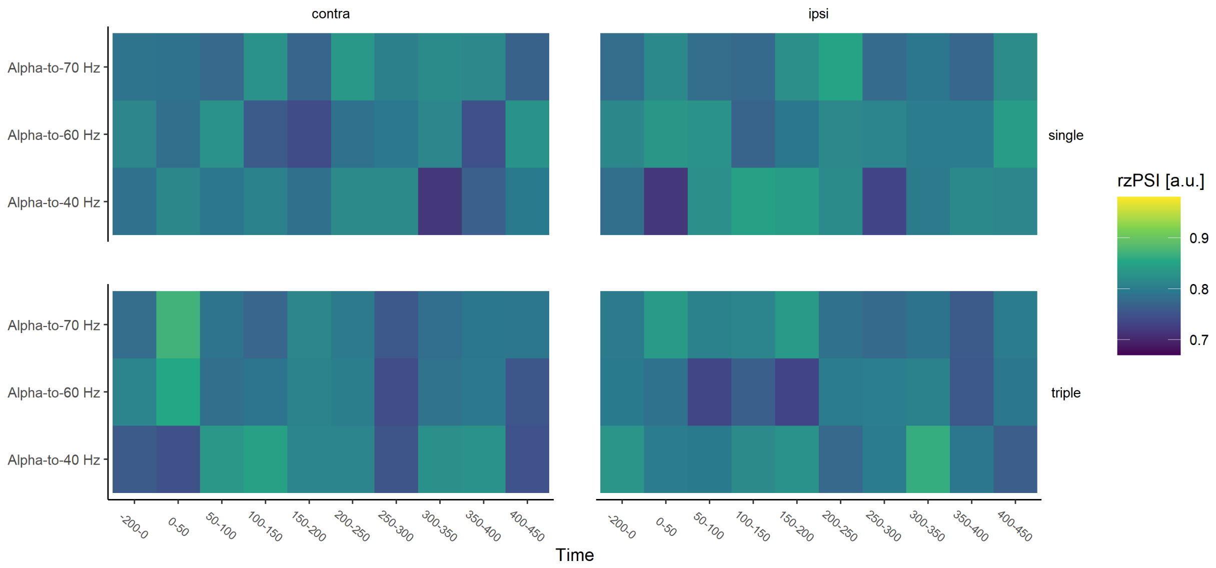

**Figure S3.3.** Cross-frequency phase synchronization indices (Rayleigh's z-transformed; rzPSIs), measuring the consistency of alpha-gamma phase difference from the right hemispheric posterior ROI in windows of 50ms length, starting at stimulus onset 0ms up to 450 ms, and in a 200ms pre-stimulus baseline, for alpha-to-40 Hz, alpha-to-60 Hz and alpha-to-70 Hz phase synchronization. Group averaged rzPSIs are shown separately for single or triple template conditions and for contralateral or ipsilateral target locations.

### S4 Control analysis: Phase locking

**Table S4.1.** Summary of model fit for theta phase locking factors (rzPLFs), from the right hemispheric ROI in the time window 150-200 ms after visual search display onset.

| Linear mixed model fit by REML | | | | |
|---|---|---|---|---|
| REML criterion at convergence: 928 | | | | |
| Scaled residuals: | | | | |
| Min | 1Q | Median | 3Q | Max |
| -2.07 | -0.57 | 0.06 | 0.59 | 2.01 |
| Random effects: | | | | |
| Groups | Term | Std.Dev. | | |
| SUBJ | (Intercept) | 16.273 | | |
| Residual | | 10.714 | | |
| Number of obs: 116, groups: SUBJ, 29. | | | | |

Fixed effects:

| | Estimate | Std. Error | t value |
|---|---|---|---|
| (Intercept) | 38 | 3.2 | 12 |
| CONDSinglevTriple | 12 | 2 | 5.8 |
| TARGContravIpsi | -1 | 2 | -0.51 |
| CONDSinglevTriple:TARGContravIpsi | 0.041 | 4 | 0.01 |

*Note.* The model includes a random-effects term for the intercept of individual subjects and the fixed effects COND (Single, Triple), TARG (Contralateral, Ipsilateral) and the interaction between them. Contrast coefficients with absolute *t* values larger than 1.96 can be treated as approximating the 5% significance level.
rzPLF = phase locking factor, Rayleigh's z-transformed.

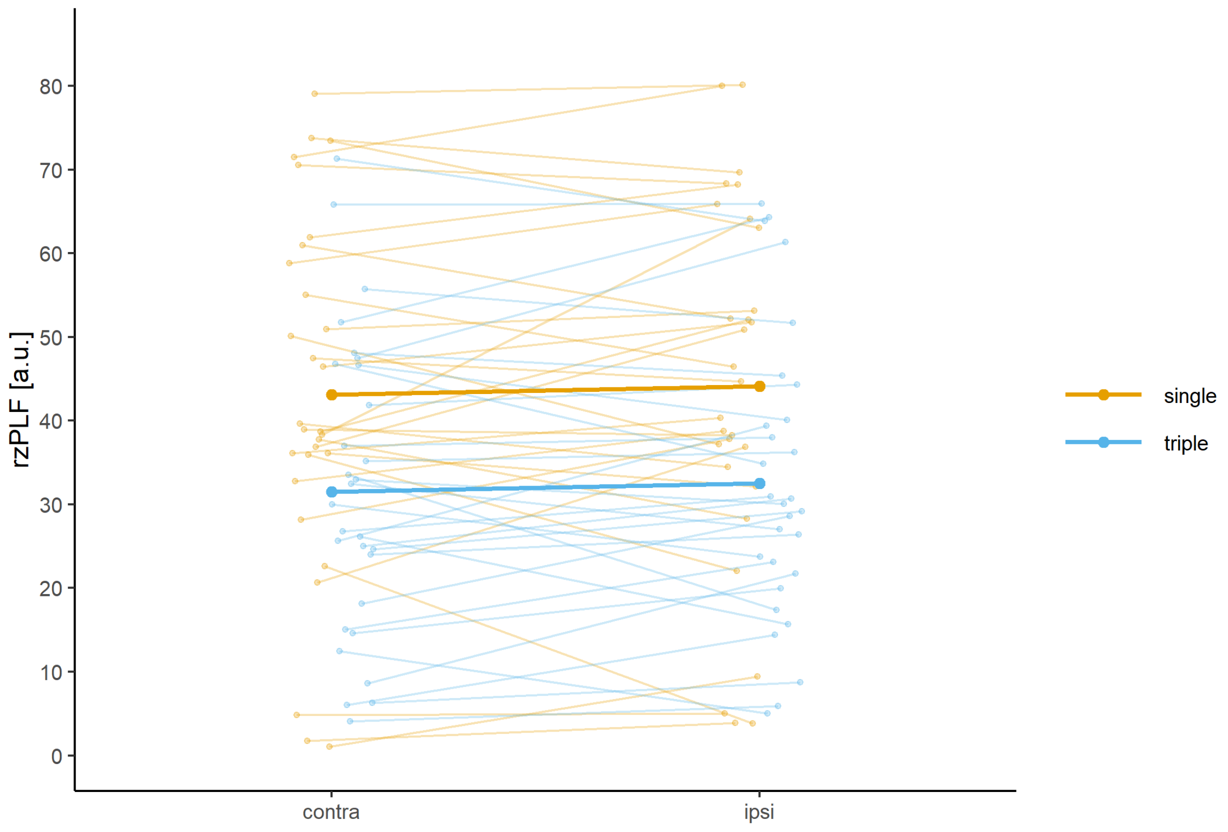

**Figure S4.1.** Theta phase locking factors (Rayleigh's z-transformed; rzPLFs), measuring the consistency of theta phase, from the right hemispheric ROI in the time window 150-200 ms after visual search display onset. RzPLFs are displayed separately for single or triple template conditions (in colour) and for contralateral or ipsilateral target locations (on the x axis). Single-subject indices (as thin lines) are overlayed with group averages (as thick lines).

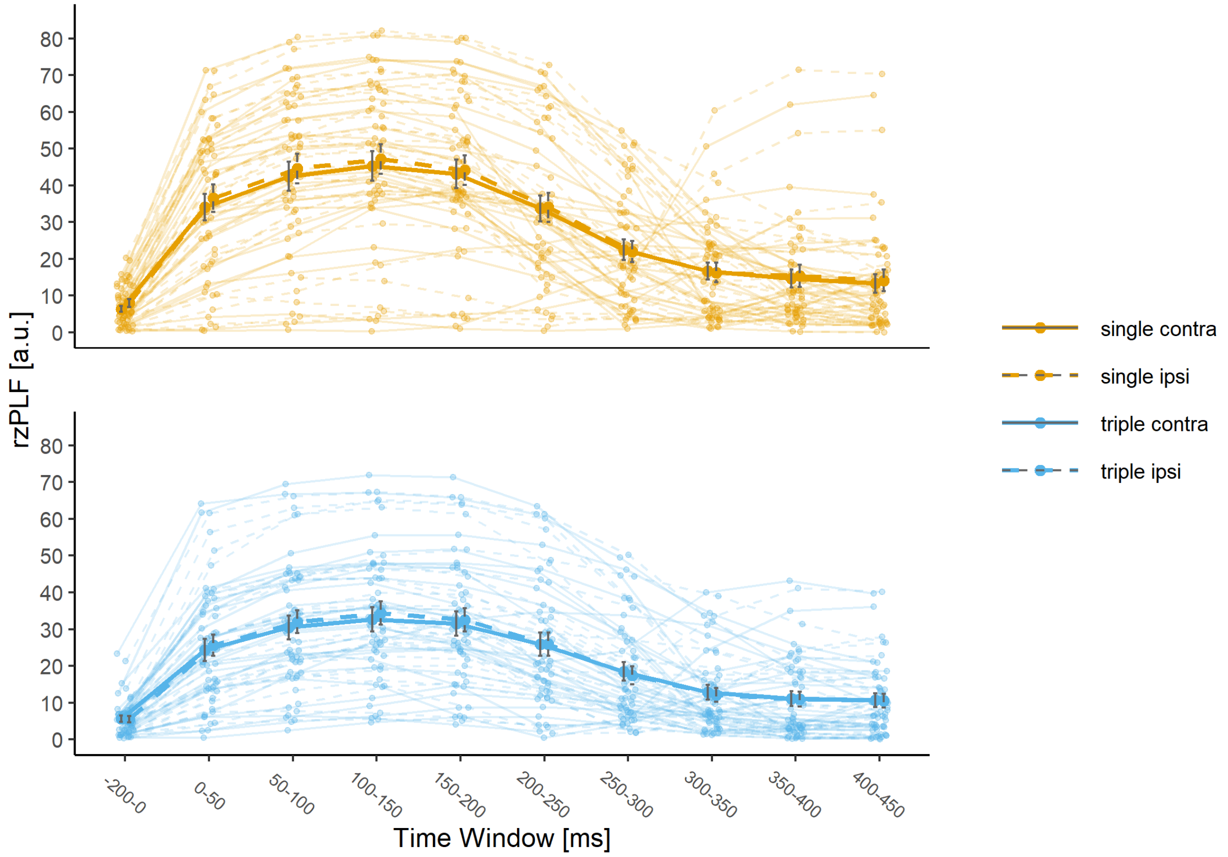

**Figure S4.2.** Theta phase locking factors (Rayleigh's z-transformed; rzPLFs), measuring the consistency of theta phase, from the right hemispheric posterior ROI in windows of 50ms length, starting at stimulus onset 0ms up to 450 ms, and in a 200ms pre-stimulus baseline. Group averaged rzPLFs are shown separately for single or triple template conditions (in color and in separate panels) and for contralateral or ipsilateral target locations (as line-type). Single-subject indices (as thin lines) are overlayed with group averages (as thick lines) and standard errors.

**Table S4.2.** Summary of model fit for gamma phase locking factors (rzPLFs), from the right hemispheric ROI in the time window 150-200 ms after visual search display onset.

| Linear mixed model fit by REML | | | | |
|---|---|---|---|---|
| REML criterion at convergence: 168 | | | | |
| Scaled residuals: | | | | |
| Min | 1Q | Median | 3Q | Max |
| -1.63 | -0.67 | -0.25 | 0.5 | 3.14 |
| Random effects: | | | | |
| Groups | Term | Std.Dev. | | |
| SUBJ | (Intercept) | 0.12818 | | |
| Residual | | 0.46565 | | |
| Number of obs: 116, groups: SUBJ, 29. | | | | |

Fixed effects:

| | Estimate | Std. Error | t value |
|---|---|---|---|
| (Intercept) | 0.95 | 0.049 | 19 |
| CONDSinglevTriple | 0.032 | 0.086 | 0.37 |
| TARGContravIpsi | -0.048 | 0.086 | -0.55 |
| CONDSinglevTriple:TARGContravIpsi | 0.15 | 0.17 | 0.89 |

*Note.* The model includes a random-effects term for the intercept of individual subjects and the fixed effects COND (Single, Triple), TARG (Contralateral, Ipsilateral) and the interaction between them. Contrast coefficients with absolute $t$ values larger than 1.96 can be treated as approximating the 5% significance level.
rzPLF = phase locking factor, Rayleigh's z-transformed.

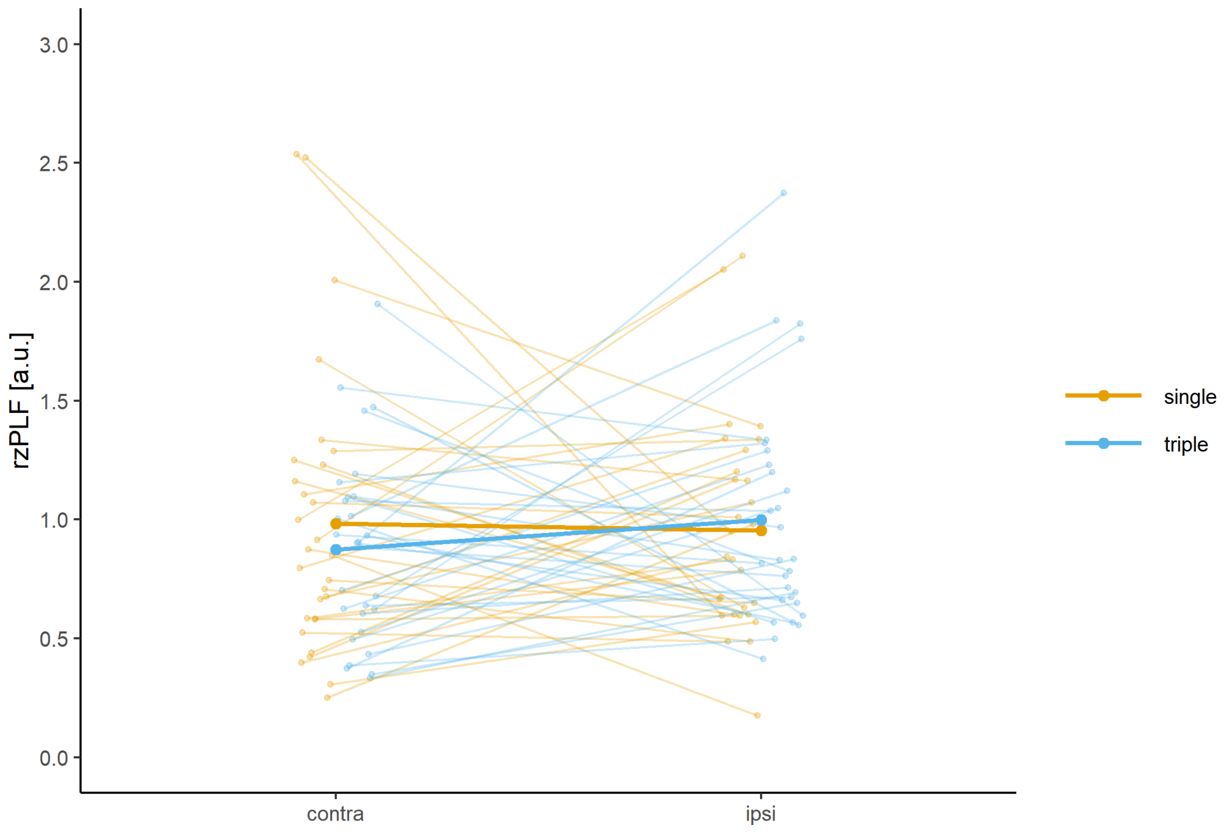

**Figure S4.3.** Gamma phase locking factors indices (Rayleigh's z-transformed; rzPLFs), measuring the consistency of gamma phase, from the right hemispheric ROI in the time window 150-200 ms after visual search display onset. RzPLFs are displayed separately for single or triple template conditions (in colour) and for contralateral or ipsilateral target locations (on the x axis) and are averaged across gammabands (center frequencies 40 Hz, 60 Hz, 70 Hz). Single-subject indices (as thin lines) are overlayed with group averages (as thick lines).

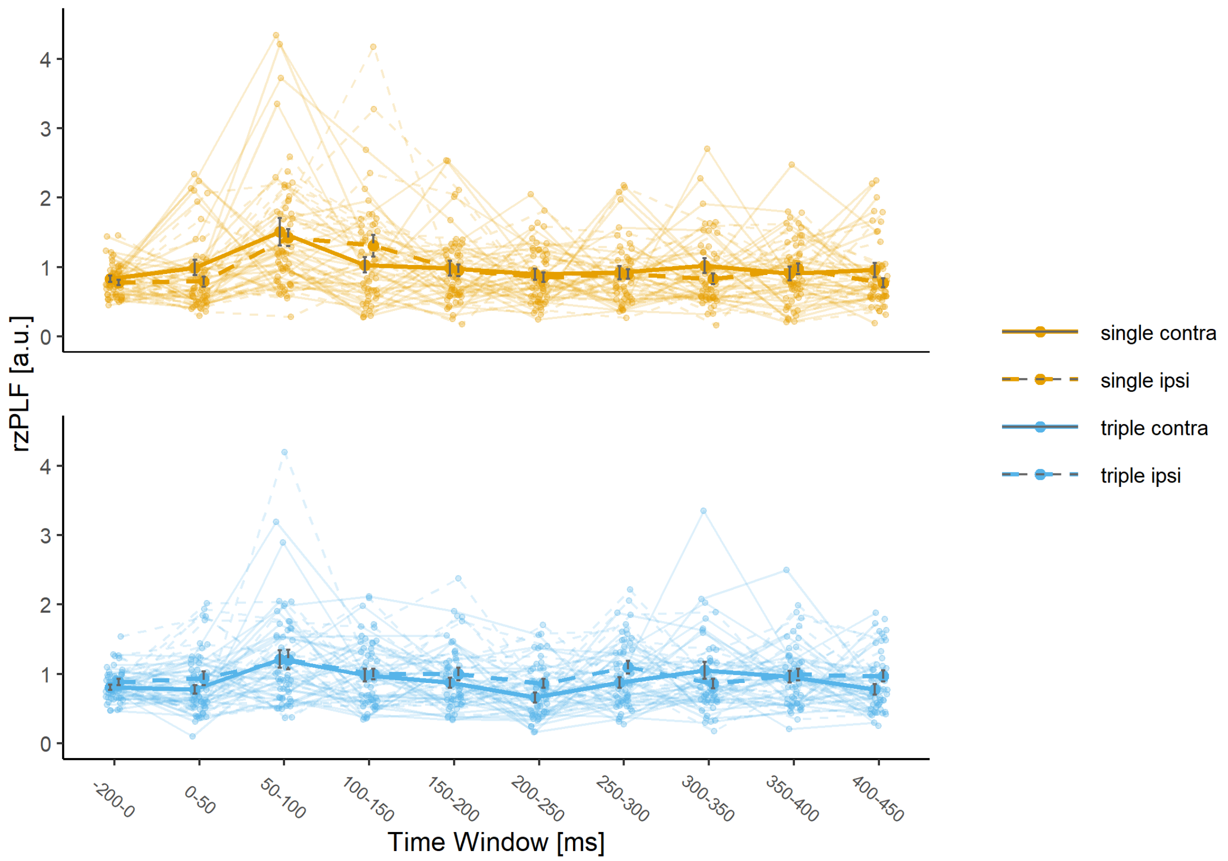

**Figure S4.4.** Gamma phase locking factors (Rayleigh's z-transformed; rzPLFs), measuring the consistency of gamma phase, from the right hemispheric posterior ROI in windows of 50ms length, starting at stimulus onset 0ms up to 450 ms, and in a 200ms pre-stimulus baseline. Group averaged rzPLFs are shown separately for single or triple template conditions (in color and in separate panels) and for contralateral or ipsilateral target locations (as line-type) and are averaged across gammabands (center frequencies 40 Hz, 60 Hz, 70 Hz). Single-subject indices (as thin lines) are overlayed with group averages (as thick lines) and standard errors.

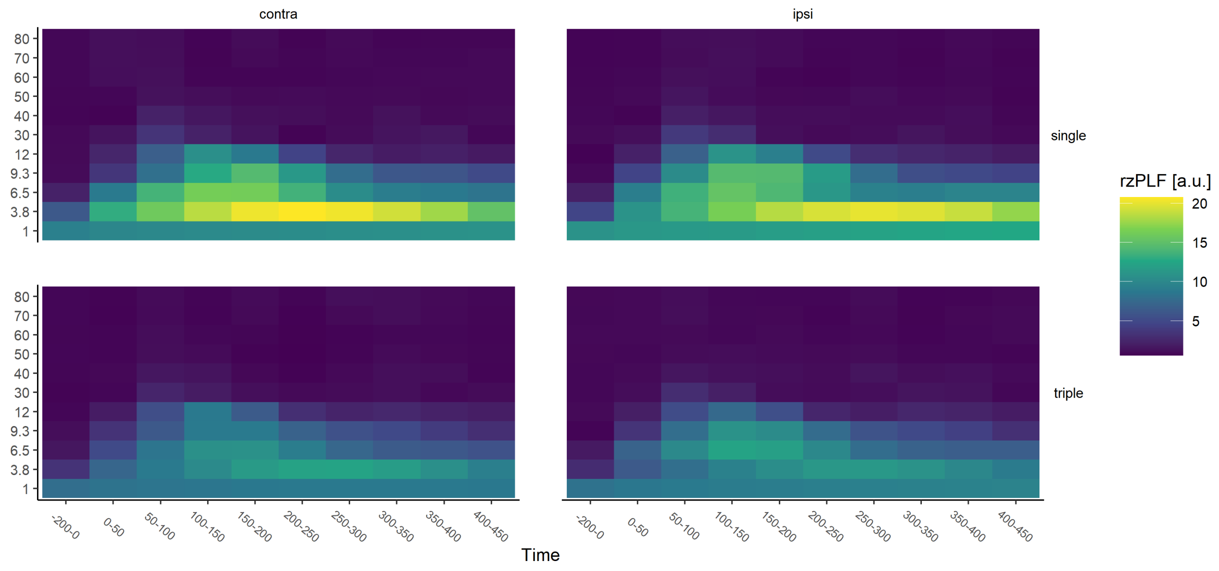

**Figure S4.5.** Phase locking factors for all extracted Morlet wavelets (Rayleigh's z-transformed; rzPLFs), measuring the consistency of phase for the extracted frequency bands (center frequencies in Hz), from the right hemispheric posterior ROI in windows of 50ms length, starting at stimulus onset 0ms up to 450 ms, and in a 200ms pre-stimulus baseline. Group averaged rzPLFs are shown separately for single or triple template conditions and for contralateral or ipsilateral target locations.

*Note.* Frequencies of interest were the theta band (center frequency 6.5 Hz) and three non-overlapping gammabands (center frequencies 40 Hz, 60 Hz, 70 Hz).

### S5 Control analysis: Amplitudes

**Table S5.1.** Summary of model fit for theta amplitudes from the right hemispheric ROI in the time window 150-200 ms after visual search display onset.

Linear mixed model fit by REML

REML criterion at convergence: 138

Scaled residuals:

| Min | 1Q | Median | 3Q | Max |
|---|---|---|---|---|
| -2.75 | -0.55 | 0 | 0.46 | 2.51 |

Random effects:

| Groups | Term | Std.Dev. |
|---|---|---|
| SUBJ | (Intercept) | 0.99267 |
| Residual | | 0.25106 |

Number of obs: 348, groups: SUBJ, 29.

Fixed effects:

| | Estimate | Std. Error | t value |
|---|---|---|---|
| (Intercept) | 2.4 | 0.19 | 13 |
| CONDSinglevTriple | 0.047 | 0.047 | 1 |
| TARGContravIpsi | 0.018 | 0.047 | 0.38 |
| CONDSinglevTriple:TARGContravIpsi | 0.062 | 0.093 | 0.66 |

*Note.* The model includes a random-effects term for the intercept of individual subjects and the fixed effects COND (Single, Triple), TARG (Contralateral, Ipsilateral) and the interaction between them. Contrast coefficients with absolute *t* values larger than 1.96 can be treated as approximating the 5% significance level.

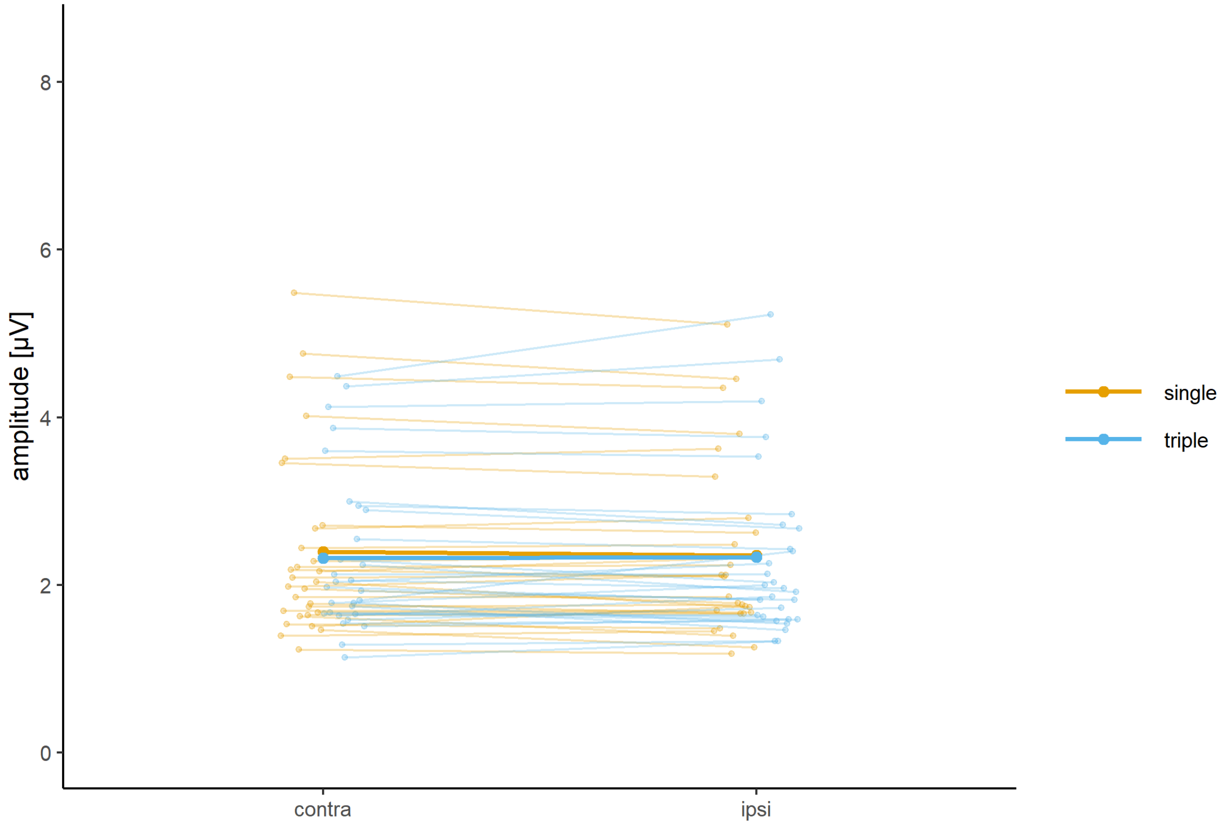

**Figure S5.1.** Theta amplitudes from the right hemispheric ROI in the time window 150-200 ms after visual search display onset. Amplitudes are displayed separately for single or triple template conditions (in colour) and for contralateral or ipsilateral target locations (on the x axis). Single-subject indices (as thin lines) are overlayed with group averages (as thick lines).

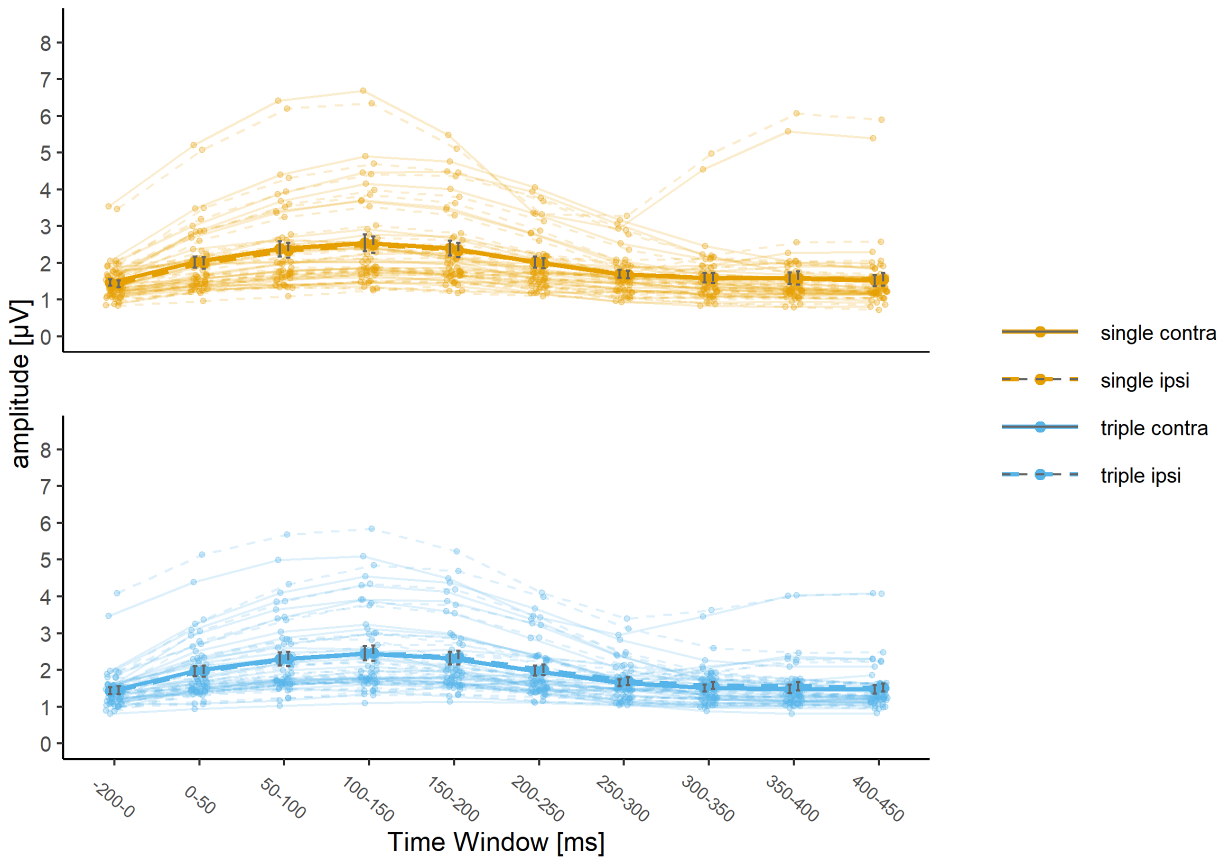

**Figure S5.2.** Theta amplitudes from the right hemispheric posterior ROI in windows of 50ms length, starting at stimulus onset 0ms up to 450 ms, and in a 200ms pre-stimulus baseline. Amplitudes are shown separately for single or triple template conditions (in colour and in separate panels) and for contralateral or ipsilateral target locations (as line-type). Single-subject indices (as thin lines) are overlayed with group averages (as thick lines) and standard errors.

**Table S5.3.** Summary of model fit for gamma amplitudes from the right hemispheric ROI in the time window 150-200 ms after visual search display onset.

Linear mixed model fit by REML

REML criterion at convergence: 250

Scaled residuals:

| Min | 1Q | Median | 3Q | Max |
|---|---|---|---|---|
| -2.6 | -0.5 | -0.03 | 0.53 | 3.39 |

Random effects:

| Groups | Term | Std.Dev. |
|---|---|---|
| SUBJ | (Intercept) | 1.53507 |
| Residual | | 0.42347 |

Number of obs: 348, groups: SUBJ, 29.

Fixed effects:

| | Estimate | Std. Error | t value |
|---|---|---|---|
| (Intercept) | 3.2 | 0.29 | 11 |
| CONDSinglevTriple | -0.068 | 0.079 | -0.86 |
| TARGContravIpsi | -0.029 | 0.079 | -0.36 |
| CONDSinglevTriple:TARGContravIpsi | 0.12 | 0.16 | 0.79 |

*Note.* The model includes a random-effects term for the intercept of individual subjects and the fixed effects COND (Single, Triple), TARG (Contralateral, Ipsilateral) and the interaction between them. Amplitudes were averaged across gammabands (center frequencies 40 Hz, 60 Hz, 70 Hz) before entering them into the model. Contrast coefficients with absolute *t* values larger than 1.96 can be treated as approximating the 5% significance level.

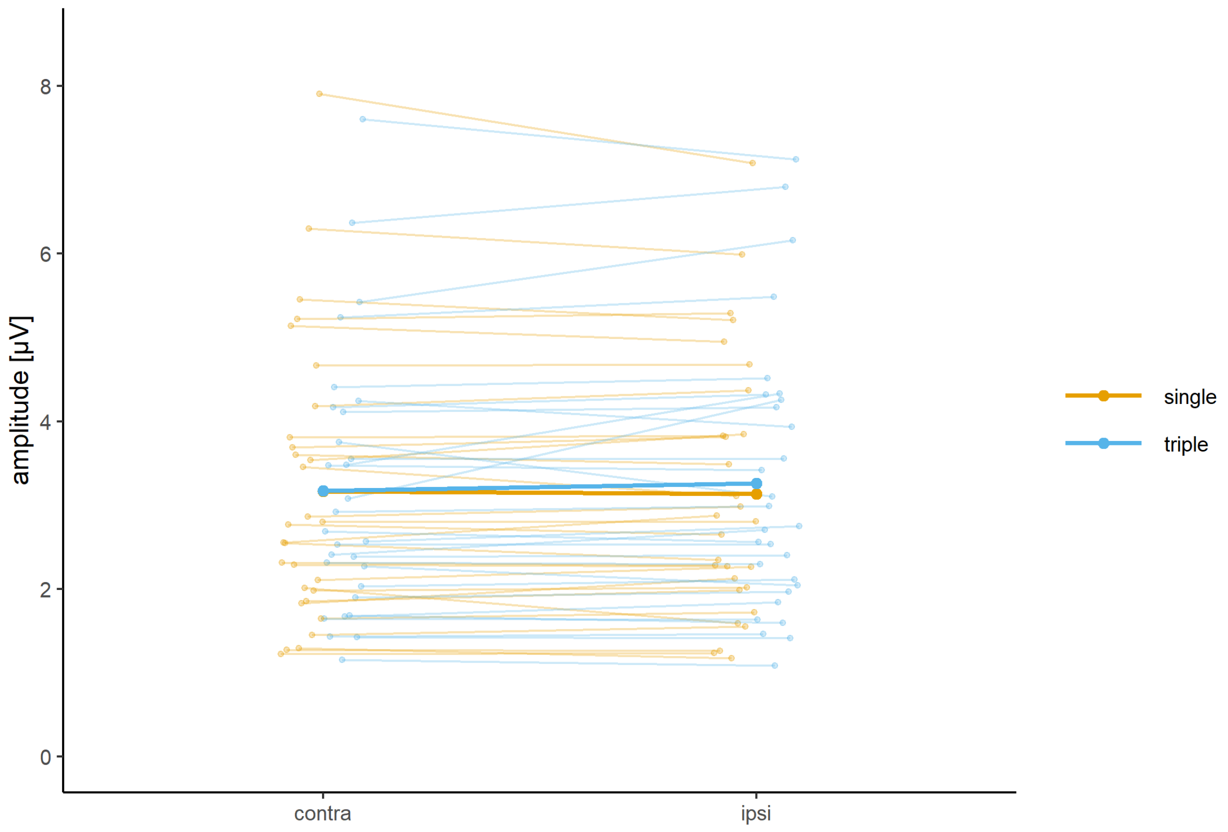

**Figure S5.3.** Gamma amplitudes from the right hemispheric ROI in the time window 150-200 ms after visual search display onset. Amplitudes are averaged across gammabands (center frequencies 40 Hz, 60 Hz, 70 Hz) and are displayed separately for single or triple template conditions (in colour) and for contralateral or ipsilateral target locations (on the x axis) and are averaged across gammabands (center frequencies 40 Hz, 60 Hz, 70 Hz). Single-subject indices (as thin lines) are overlayed with group averages (as thick lines).

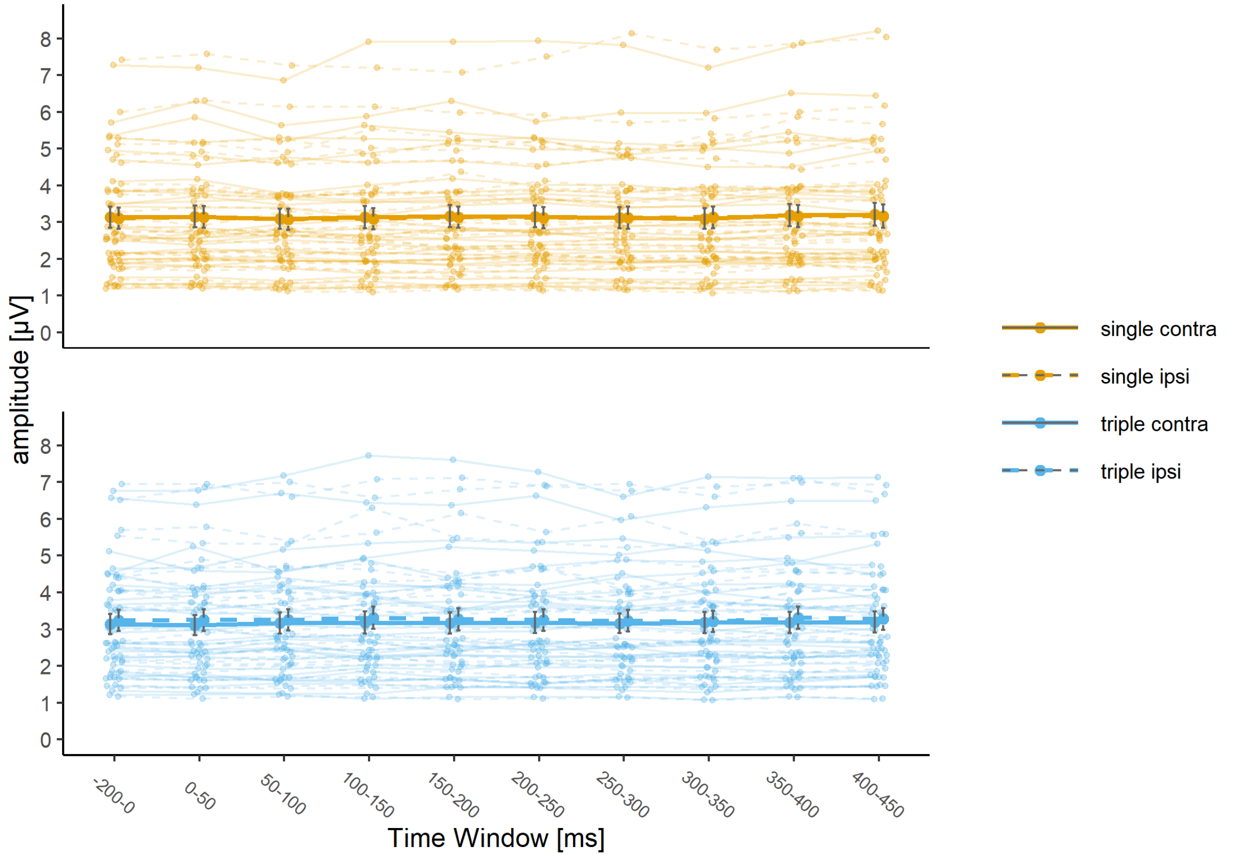

**Figure S5.4.** Gamma amplitudes from the right hemispheric posterior ROI in windows of 50ms length, starting at stimulus onset 0ms up to 450 ms, and in a 200ms pre-stimulus baseline. Amplitudes are averaged across gammabands (center frequencies 40 Hz, 60 Hz, 70 Hz) and are shown separately for single or triple template conditions (in colour and in separate panels) and for contralateral or ipsilateral target locations (as line-type). Single-subject indices (as thin lines) are overlayed with group averages (as thick lines) and standard errors.

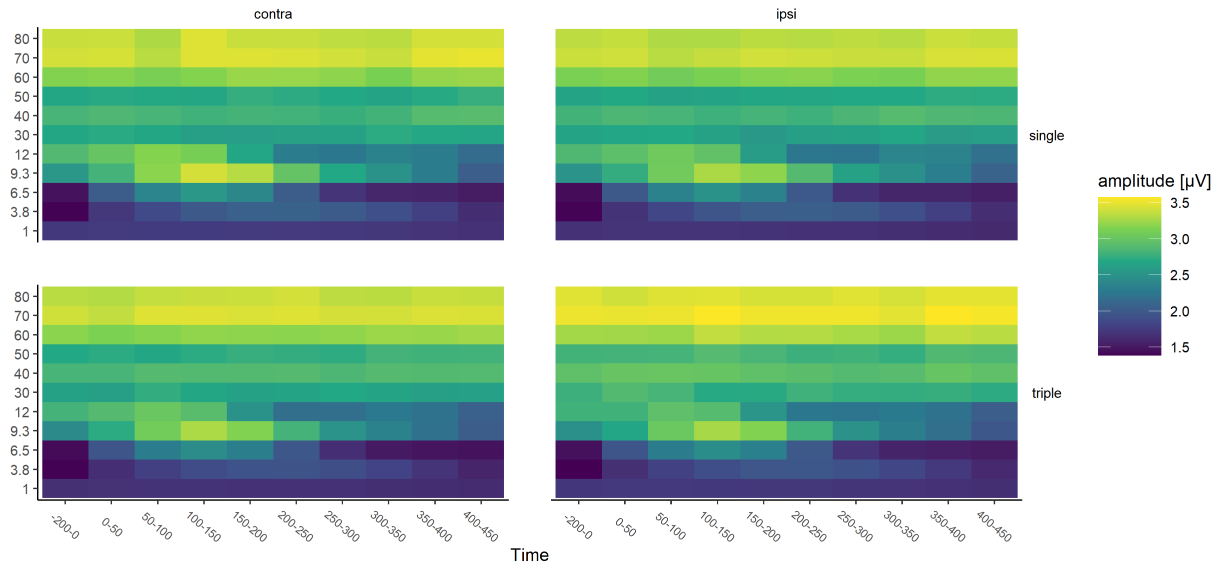

**Figure S5.5.** Amplitudes for all extracted Morlet wavelets (center frequencies in Hz), from the right hemispheric posterior ROI in windows of 50ms length, starting at stimulus onset 0ms up to 450 ms, and in a 200ms pre-stimulus baseline. Group averaged amplitudes are shown separately for single or triple template conditions and for contralateral or ipsilateral target locations.
*Note.* Frequencies of interest were the theta band (center frequency 6.5 Hz) and three non-overlapping gammabands (center frequencies 40 Hz, 60 Hz, 70 Hz).

### S6 Control analysis based on data with matched number of trials

To account for the number of trials (n) that went into calculation of the phase synchonization index, PSI values were transformed using Rayleigh's z transform since usually, measures of phase-synchronization are sensitive to the difference of trial numbers across conditions used (Cohen, 2014). Naturally, this does not eliminate the difference in signal-to-noise ratio between conditions, which was most likely lower in the triple template condition. Therefore, the goal of this additional control analysis was to analyse a dataset where trial numbers are equalized between single- and triple-template conditions, such that we draw a random subset of the same number of trials in the condition with fewer trials.

In the main analysis, we had used all correct and artifact-free trials. In this additional control analysis, we sampled from the template condition with a larger pool of trials, while using all trials from the template condition with fewer trials. We wanted to keep as many trials as possible in the analysis and since there was no systematic difference in trials for targets presented in the left or right hemifield, this sampling procedure was administered separately for each targets location. First, we determined the minimal number of trials in the single or triple template condition, separately for targets presented at the left and right hemifield. Then, for each target location, their minimal trial number determined the size of the random subset of trials that was drawn from the larger pool of trials. Based on this data equalized by trial amount, PSIs were calculated across an average of 51 trials (SD= 11.2) for left hemifield targets and 53 trials (SD=13.2) for right hemifield targets. All other analysis septs were identical to the main analysis. We report results for the right-hemispheric ROI.

Importantly, results from this control analysis are very similar to those obtained from the main analysis based on all trials. The critical interaction between COND and TARG from the main analysis (0.18, $t = 3.6$, see figure 4B) was reproduced for the analysis on trial-matched data

(0.11, t = 2.1), which shows the same pattern of results that the single template condition showed larger estimates for contra compared to ipsilateral targets, whereas this was not the case for the triple condition. Like in the main analysis, where rzPSIS for Theta-to-70Hz were smaller than for Theta-to-60Hz (-0.068, t = -2.2) and this gamma frequency dependent difference in rzPSIs was smaller for single than triple template conditions (0.15, t = 2.5, see figure 4C), in the control analysis with the trial-matched data, a similar main effect for CFSTheta70vTheta60 (-0.077, t=-2.4) and a similar interaction with COND (0.15, t=2.3) reproduce this pattern of results. All other effects were not substantial (t values below 1.96).

**Table S6.1.** Summary of model fit for trial-matched theta-gamma phase synchronization indices (rzPSIs) from the right hemispheric ROI in the time window 150-200 ms after visual search display onset.

Linear mixed model fit by REML

REML criterion at convergence: 61

Scaled residuals:

| Min | 1Q | Median | 3Q | Max |
|---|---|---|---|---|
| -2.52 | -0.66 | -0.01 | 0.54 | 4.06 |

Random effects:

| Groups | Term | Std.Dev. |
|---|---|---|
| SUBJ | (Intercept) | 0.044999 |
| Residual | | 0.245836 |

Number of obs: 348, groups: SUBJ, 29.

Fixed effects:

| | | | |
|---|---|---|---|
| (Intercept) | 0.82 | 0.016 | 52 |
| CONDSinglevTriple | -0.0058 | 0.026 | -0.22 |
| TARGContravIpsi | 0.023 | 0.026 | 0.86 |
| CFSTheta60vTheta40 | -0.036 | 0.032 | -1.1 |
| CFSTheta70vTheta60 | -0.077 | 0.032 | -2.4 |
| CONDSinglevTriple:TARGContravIpsi | 0.11 | 0.053 | 2.1 |
| CONDSinglevTriple:CFSTheta60vTheta40 | -0.12 | 0.065 | -1.8 |
| CONDSinglevTriple:CFSTheta70vTheta60 | 0.15 | 0.065 | 2.3 |
| TARGContravIpsi:CFSTheta60vTheta40 | 0.056 | 0.065 | 0.86 |
| TARGContravIpsi:CFSTheta70vTheta60 | -0.074 | 0.065 | -1.1 |
| CONDSinglevTriple:TARGContravIpsi:CFSTheta60vTheta40 | -0.046 | 0.13 | -0.36 |
| CONDSinglevTriple:TARGContravIpsi:CFSTheta70vTheta60 | 0.0025 | 0.13 | 0.019 |

*Note.* The model includes a random-effects term for the intercept of individual subjects and the fixed effects COND (Single, Triple), TARG (Contralateral, Ipsilateral), CFS (Theta-to-40 Hz, Theta-to-60 Hz, Theta-to-70 Hz), and interactions between them. Contrast coefficients with absolute *t* values larger than 1.96 can be treated as approximating the 5% significance level. rzPSI = cross-frequency phase synchronization index, Rayleigh's z-transformed.

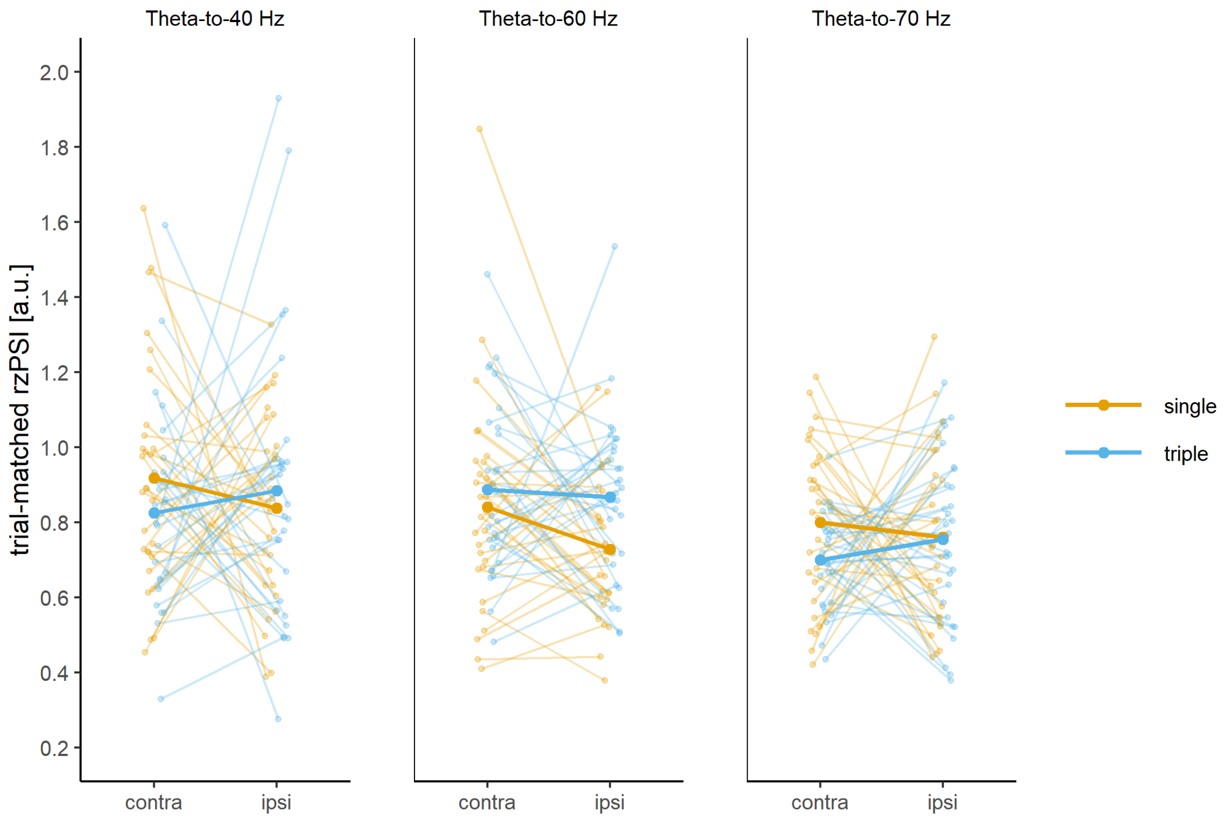

**Figure S6.1.** Trial-matched cross-frequency phase synchronization indices (Rayleigh's z-transformed; rzPSIs), measuring the consistency of theta-gamma phase difference based on a dataset with the same number of trials in both template conditions, from the right hemispheric ROI in the time window 150-200 ms after visual search display onset. Trial-matched RzPSIs are displayed separately for single or triple template conditions (in colour), for contralateral or ipsilateral target locations (on the x axis) and for theta-to-40 Hz, theta-to-60 Hz or theta-to-70 Hz cross-frequency synchronization (in separate panels). Single-subject indices (as thin lines) are overlayed with group averages (as thick lines).

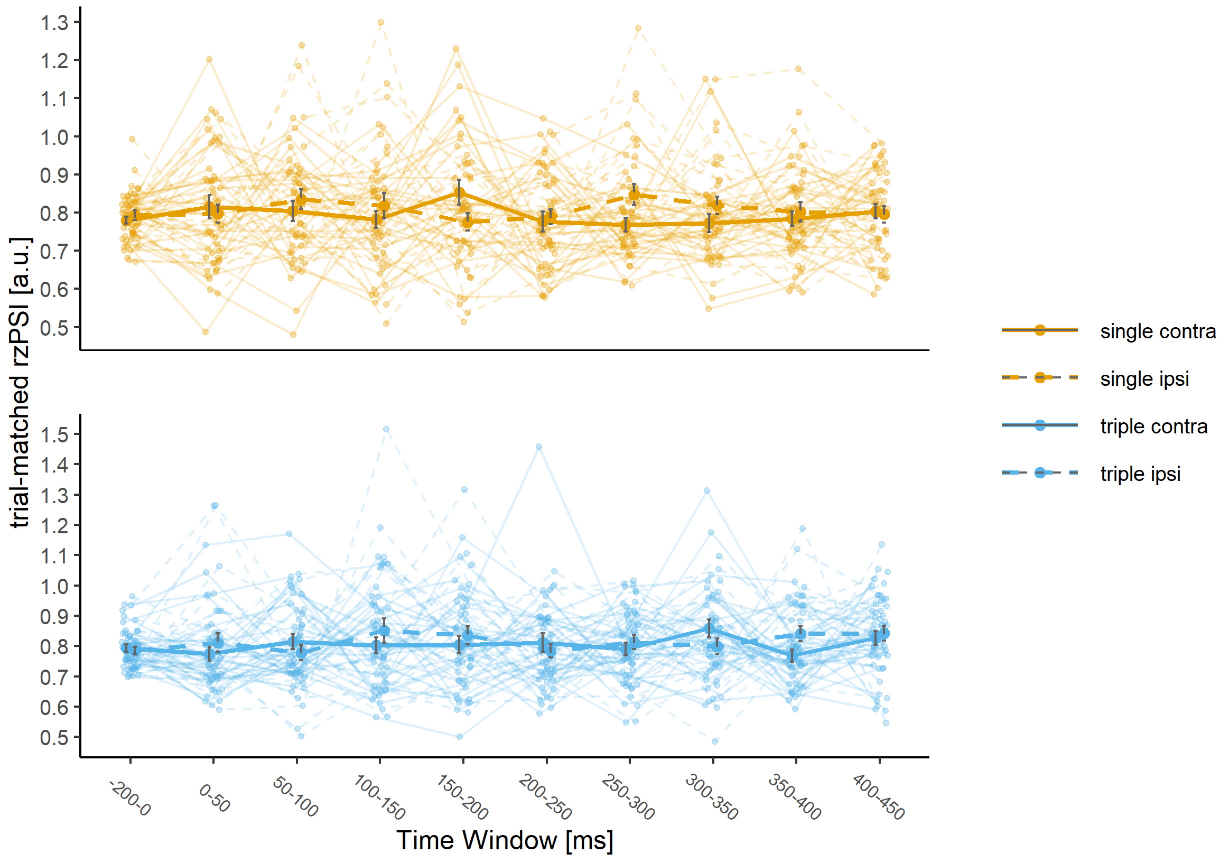

**Figure S6.2.** Trial-matched cross-frequency phase synchronization indices (Rayleigh's z-transformed; rzPSIs), measuring the consistency of theta-gamma phase difference based on a dataset with the same number of trials in both template conditions, from the right hemispheric posterior ROI in windows of 50ms length, starting at stimulus onset 0ms up to 450 ms, and in a 200ms pre-stimulus baseline. Group averaged trial-matched rzPSIs are shown separately for single or triple template conditions (in color and in separate panels) and for contralateral or ipsilateral target locations (as line-type). Indices are averaged across theta-to-40 Hz, theta-to-60 Hz or theta-to-70 Hz cross-frequency synchronization. Single-subject indices (as thin lines) are overlayed with group averages (as thick lines) and standard errors.

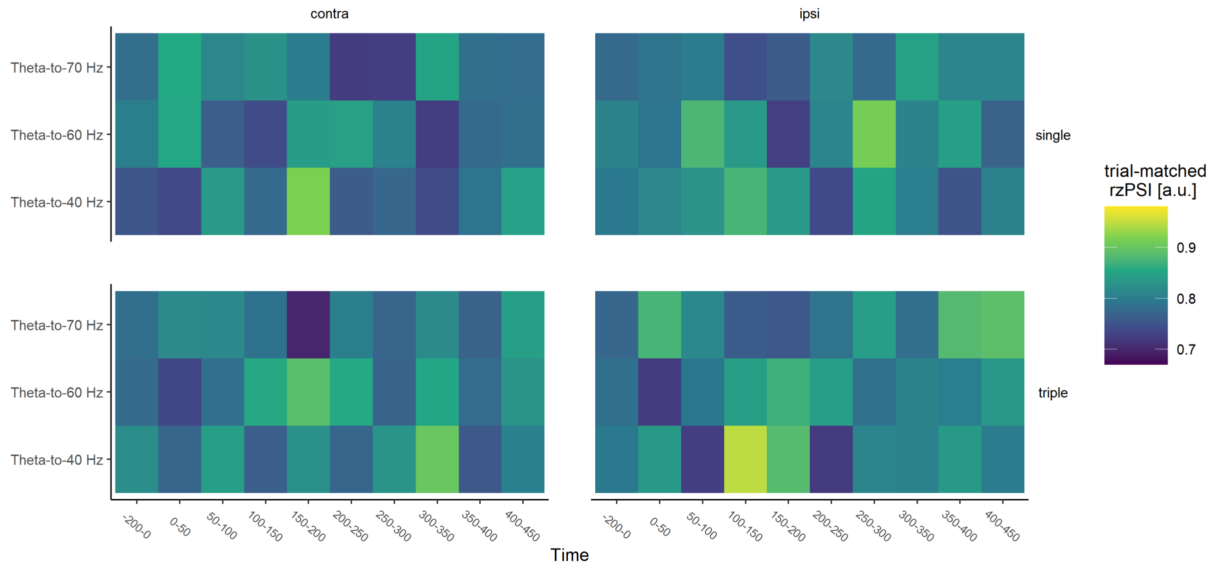

**Figure S6.3.** Trial-matched cross-frequency phase synchronization indices (Rayleigh's z-transformed; rzPSIs), measuring the consistency of theta-gamma phase difference based on a dataset with the same number of trials in both template conditions, from the right hemispheric posterior ROI in windows of 50ms length, starting at stimulus onset 0ms up to 450 ms, and in a 200ms pre-stimulus baseline, for theta-to-40 Hz, theta-to-60 Hz and theta-to-70 Hz phase synchronization. Group averaged trial-matched rzPSIs are shown separately for single or triple template conditions and for contralateral or ipsilateral target locations.

### S7 Control analysis based on surrogate data

**Table S7.1.** Summary of model fit for trial-shuffled theta-gamma phase synchronization indices (rzPSIs) from the right hemispheric ROI in the time window 150-200 ms after search onset.

| Linear mixed model fit by REML | | | |
|---|---|---|---|
| REML criterion at convergence: 13 | | | |
| Scaled residuals: | | | |
| Min 1Q Median 3Q Max | | | |
| -2.99 -0.63 -0.04 0.54 4.38 | | | |
| Random effects: | | | |
| Groups Term Std.Dev. | | | |
| SUBJ (Intercept) 0.00000 | | | |
| Residual 0.23217 | | | |
| Number of obs: 348, groups: SUBJ, 29. | | | |
| Fixed effects: | | | |
| (Intercept) | 0.84 | 0.012 | 67 |
| CONDSinglevTriple | -0.03 | 0.025 | -1.2 |
| TARGContravIpsi | -0.012 | 0.025 | -0.48 |
| CFSTheta60vTheta40 | -0.087 | 0.03 | -2.9 |
| CFSTheta70vTheta60 | 0.015 | 0.03 | 0.49 |
| CONDSinglevTriple:TARGContravIpsi | -0.055 | 0.05 | -1.1 |
| CONDSinglevTriple:CFSTheta60vTheta40 | 0.048 | 0.061 | 0.79 |
| CONDSinglevTriple:CFSTheta70vTheta60 | -0.12 | 0.061 | -1.9 |
| TARGContravIpsi:CFSTheta60vTheta40 | -0.053 | 0.061 | -0.86 |
| TARGContravIpsi:CFSTheta70vTheta60 | 0.028 | 0.061 | 0.46 |
| CONDSinglevTriple:TARGContravIpsi:CFSTheta60vTheta40 | -0.31 | 0.12 | -2.5 |
| CONDSinglevTriple:TARGContravIpsi:CFSTheta70vTheta60 | 0.054 | 0.12 | 0.44 |
| (Intercept) | 0.84 | 0.012 | 67 |

*Note.* The model includes a random-effects term for the intercept of individual subjects and the fixed effects COND (Single, Triple), TARG (Contralateral, Ipsilateral), CFS (Theta-to-40 Hz, Theta-to-60 Hz, Theta-to-70 Hz), and interactions between them. Contrast coefficients with absolute $t$ values larger than 1.96 can be treated as approximating the 5% significance level. rzPSI = cross-frequency phase synchronization index, Rayleigh's z-transformed.

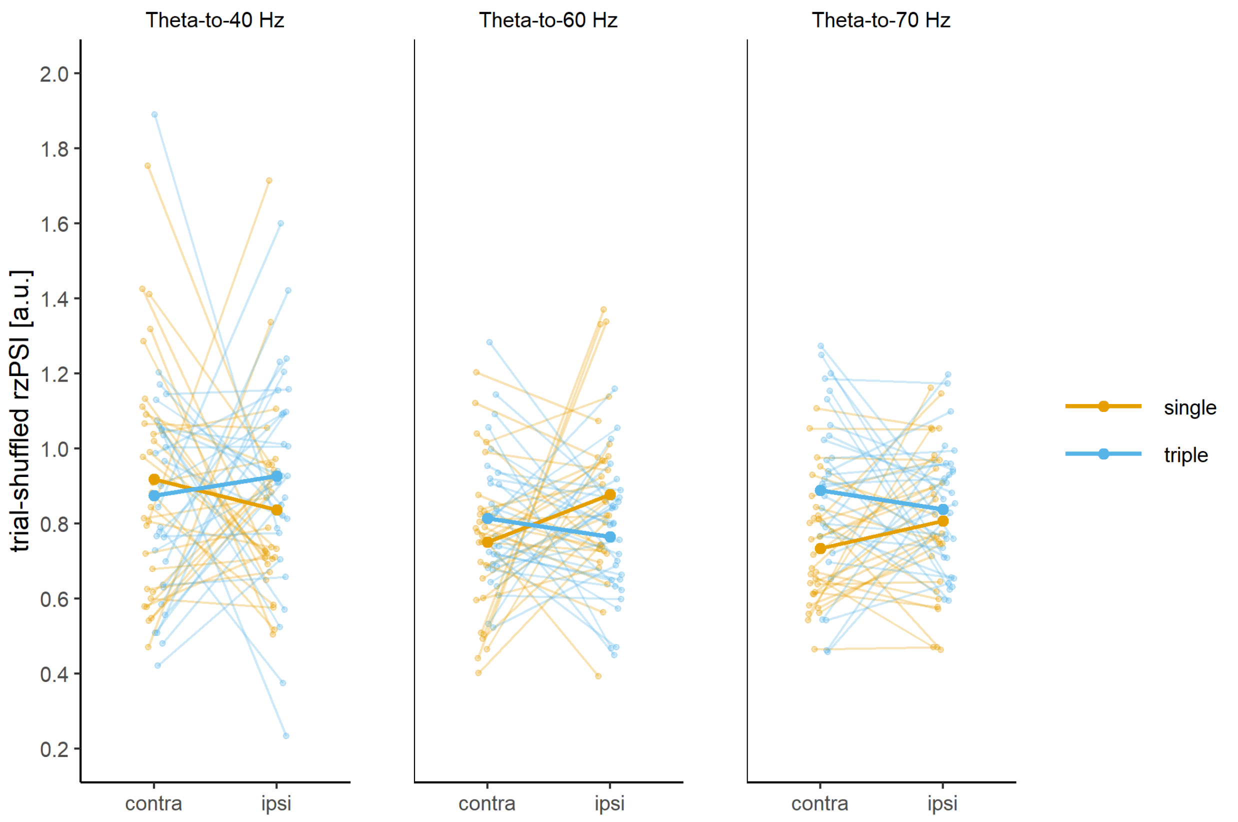

**Figure S7.1.** Trial-shuffled cross-frequency phase synchronization indices (Rayleigh's z-transformed; rzPSIs), measuring the consistency of theta-gamma phase difference based on a surrogate dataset, from the right hemispheric ROI in the time window 150-200 ms after visual search display onset. Trial-shuffled RzPSIs are displayed separately for single or triple template conditions (in colour), for contralateral or ipsilateral target locations (on the x axis) and for theta-to-40 Hz, theta-to-60 Hz or theta-to-70 Hz cross-frequency synchronization (in separate panels). Single-subject indices (as thin lines) are overlayed with group averages (as thick lines).

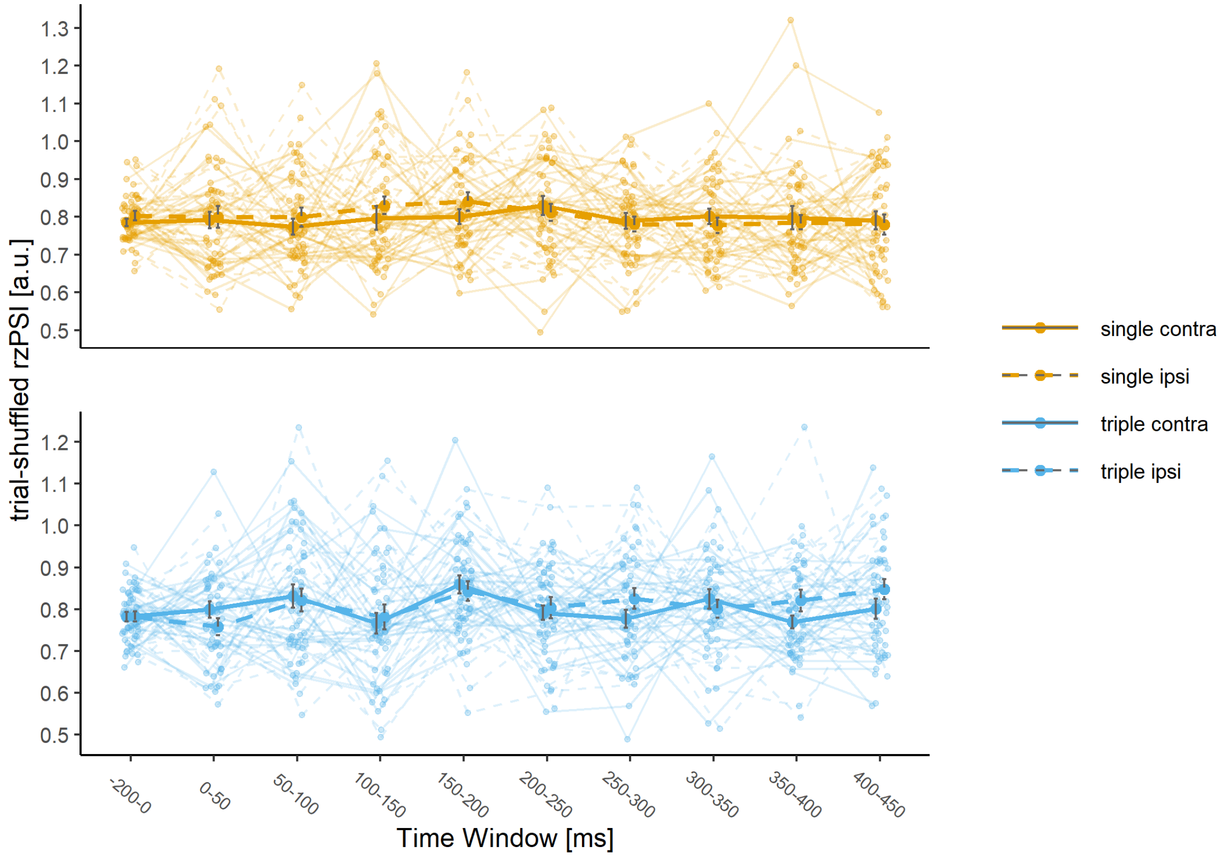

**Figure S7.2.** Trial-shuffled cross-frequency phase synchronization indices (Rayleigh's z-transformed; rzPSIs), measuring the consistency of theta-gamma phase difference based on a surrogate dataset, from the right hemispheric posterior ROI in windows of 50ms length, starting at stimulus onset 0ms up to 450 ms, and in a 200ms pre-stimulus baseline. Group averaged trial-matched rzPSIs are shown separately for single or triple template conditions (in color and in separate panels) and for contralateral or ipsilateral target locations (as line-type). Indices are averaged across theta-to-40 Hz, theta-to-60 Hz or theta-to-70 Hz cross-frequency synchronization. Single-subject indices (as thin lines) are overlayed with group averages (as thick lines) and standard errors.

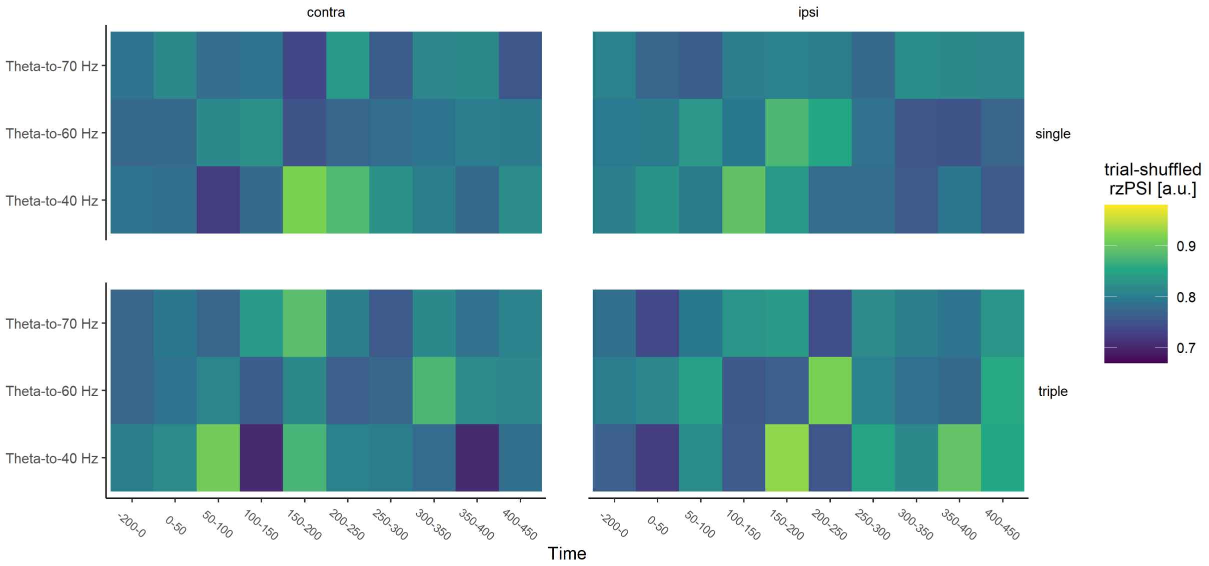

**Figure S6.3.** Trial-shuffled cross-frequency phase synchronization indices (Rayleigh's z-transformed; rzPSIs), measuring the consistency of theta-gamma phase difference based on a surrogate dataset, from the right hemispheric posterior ROI in windows of 50ms length, starting at stimulus onset 0ms up to 450 ms, and in a 200ms pre-stimulus baseline, for theta-to-40 Hz, theta-to-60 Hz and theta-to-70 Hz phase synchronization. Group averaged trial-shuffled rzPSIs are shown separately for single or triple template conditions and for contralateral or ipsilateral target locations.

### S8 Is there a relationship to task accuracy or to N2pc amplitude?

We investigated whether response accuracy was correlated with the observed rzPSIs from the right hemispheric ROI in the time window 150-200 ms, separately for Theta-to-40Hz, Theta-to-60 Hz and Theta-to-70 Hz CFS and for both template conditions and for targets presented in the left (contralateral) or right (ipsilateral) hemifield. If rzPSIs were predictive of response accuracy, we would expect to see a significant correlation. This was not the case: All correlation coefficients did not exceed the significance threshold (all ps > .05). Please see figure S8.1 for a visualization.

Next, we investigated whether the N2pc amplitude was correlated with the observed rzPSIs from the right hemispheric ROI in the time window 150-200 ms. We computed the average N2pc amplitude in the time window 200-350 ms for the difference between contra minus ipsilateral sites relative to target location. Average N2pc amplitudes significantly differed between the template conditions ($t(28)=-3.6$, $p=0.001$, paired-samples t-test). Because the N2pc is calculated for contralateral minus ipsilateral sites, we also built a difference value for rzPSIs by subtracting ipsilateral from contralateral rzPSIs. Again, we would expect to see a significant correlation if rzPSIs were predictive of N2pc amplitudes. This was not the case: none of the correlation coefficients for either combination of template conditions and Theta-to-40Hz, Theta-to-60 Hz and Theta-to-70 Hz CFS exceeded the significance threshold (all ps > .05). Please see figure S8.2 for a visualization.

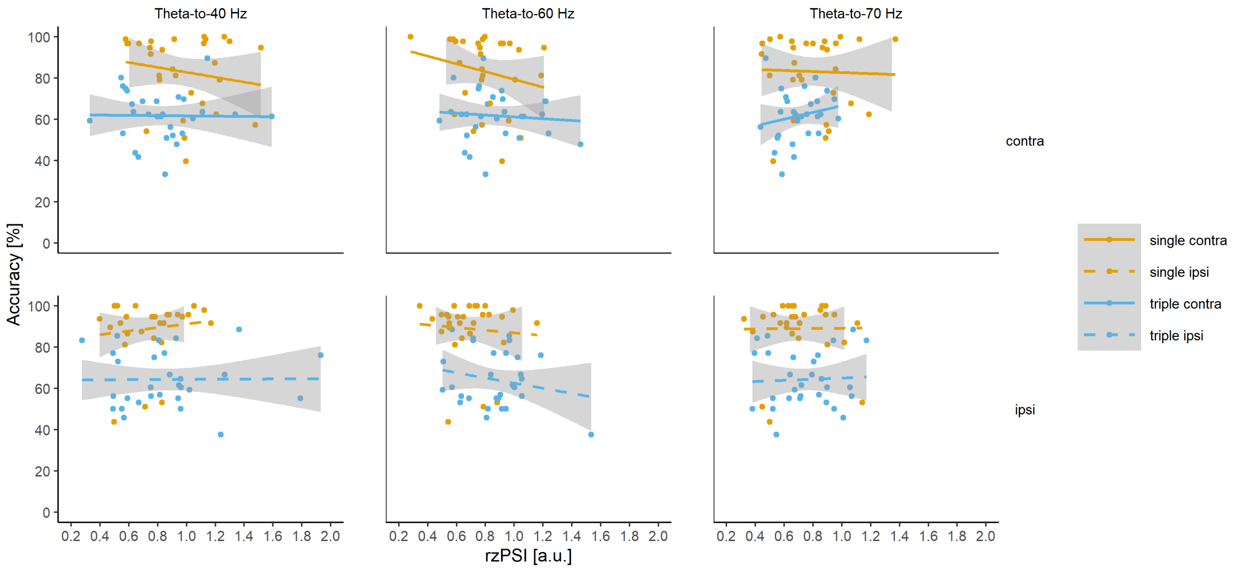

**Figure S8.1.** Relationship between task accuracy and cross-frequency phase synchronization indices (Rayleigh's z-transformed; rzPSIs) from the right hemispheric ROI in the time window 150-200 ms after visual search display onset. Data are displayed separately for single or triple template conditions (in colour), for contralateral or ipsilateral target locations (in separate panels) and for theta-to-40 Hz, theta-to-60 Hz or theta-to-70 Hz cross-frequency synchronization (in separate panels).

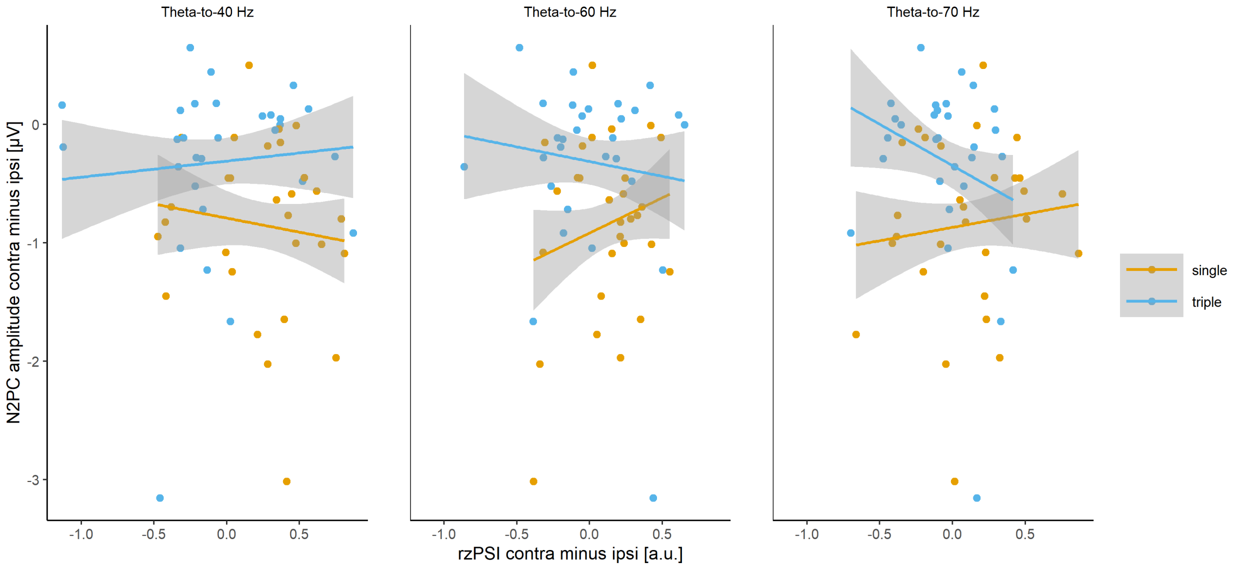

**Figure S8.2.** Relationship between N2pc amplitudes and contra minus ipsilateral cross-frequency phase synchronization indices (Rayleigh's z-transformed; rzPSIs) from the right hemispheric ROI in the time window 150-200 ms after visual search display onset. Data are displayed separately for single or triple template conditions (in colour) and for theta-to-40 Hz, theta-to-60 Hz or theta-to-70 Hz cross-frequency synchronization (in separate panels).

### S9 Are there target switch costs in the triple template condition?

Interestingly, the assumption that templates are being matched sequentially makes direct predictions about switch costs between trials where a different or the same target as on the previous trial is presented. Because the paradigm was not designed to contrast stay and switch trials, there is a much lower number of stay trials than switch trials. After excluding trials with incorrect responses and artifacts, this does not leave us with a sufficient number of trials to obtain reliable PSI measures for both types of trials. We can only speculate about how PSIs might behave in a different experimental design which would be designed explicitly to contrast stay and switch trials. Based on studies using oscillatory analyses to analyze switch costs in visual search (deVries et al., 2018; van Driel et al., 2019) and based on evidence from oscillatory analyses on more classical task switching paradigms (Capizzi et al., 2020; Cooper et al., 2019; McKewen et al., 2020), one would expect that especially neuronal networks over frontal cortical regions, oscillating at slow frequencies in would play a key role in task switching, however, these are not reflected in our source model. In our current work and in previous studies that we build on (Sauseng et al., 2008 and Holz et al., 2010), we observed the CSF effects for posterior regions of interest. For posterior regions, there is some evidence for switch costs in event-related components evoked by a visual search task, as reflected by longer latencies for N2pc components (Becker et al., 2014; Töllner et al., 2008; Rangelov et al., 2013) or for MVPA decoding performance in the N2pc time window (Ort et al., 2019) following switches, but it is unclear, how exactly switch costs may be reflected in oscillatory phase-coupling measures here. However, we think that the possible scenarios of obtaining differences in PSIs between a multiple template condition compared to a single template condition could inform a hypothesis about the expected results when comparing switch trials in a multiple template condition to stay trials in a multiple template condition. Overall, this is speculative and we do not wish to make strong predictions here.

We initially hypothesized that in the condition where one out of multiple targets must be found, the timing of the matching process would exhibit more temporal variability than for a single possible target. This predicts that cross-frequency phase relations vary over trials, such that low estimates will be achieved. So for a different experimental design which would be designed explicitly to investigate switch costs, one possibility would be to expect a similar scenario for a contrast between switch and stay trials, such that lower indices would be expected for switch trials. In the light of our results (where we observed such an effect for a contrast between triple and single conditions at the right-hemispheric ROI in the TOI) and given that in our design, the triple condition contains a majority of switch trials, this seems rather plausible. Yet, we would still expect that stay trials in a multiple templates condition show a difference to a single template condition because only in the single template case, we can expect a temporally very precise matching across trials.

Another possibility, however, would be to expect that when one out of multiple targets must be found, the matching process would happen consistently later, but with low temporal variability across trials. In this case, we would have expected to observe a later effect in the triple condition, but with high estimates of phase synchrony, comparable to those in the single condition. Although this was not the case for this contrast in our current results, such a pattern may be expected for a contrast between switch vs. stay trials. However, hybrid versions could be possible, for example, if memory matching in switch trials happened with great temporal variability and also consistently later; then, low estimates would be expected as well.

However, while these predictions are straightforward for experimental paradigms that allow a precise measurement of reaction times, they are less clear for our experimental paradigm, both for behavioural and brain-based measures. Due to the rather complex stimuli and short display times, the task in our study, especially the triple condition, was quite challenging for subjects and so our instructions stressed the importance of accurate over fast responses. Therefore, response times measured here are not speeded reaction times in the classical sense. This is particularly true for the triple template condition when compared to the single template condition, we observed a substantial slowing in button presses as well as an increase in variance. With this in mind, response times must be interpreted with caution and are, by design, not sensitive enough to reflect costs of switching on a trial-by-trial basis. Concerning the EEG-based dependent variable we analyze, these PSIs would also not be expected to be sensitive to switch costs. Inherently in the experimental design, there are switch and stay trials, however, there is a much lower number of stay trials than switch trials because the paradigm was not designed to contrast stay and switch trials. After excluding trials with incorrect responses and artifacts, this does not leave us with a sufficient number of trials to obtain reliable PSI measures for both types of trials.
